# ABA-induced SA accumulation causes higher susceptibility in *B. juncea* as compared to tolerant genotypes against *A. brassicae*

**DOI:** 10.1101/2022.04.28.489833

**Authors:** Shikha Dixit, Anita Grover, Suchitra Pushkar, Shashi Bala Singh

**Author notes:** Corresponding Author; Dr. Shikha Dixit National Institute for Plant Biotechnology, Lal Bahadur Shastri Building, Pusa Campus, New Delhi, India-110012. Affiliation(s) and address (es) of the author(s): 1. Dr. Shikha Dixit (corresponding author) National Institute for Plant Biotechnology, Lal Bahadur Shastri Building, Pusa Campus, New Delhi, India-110012 2. Dr. Anita Grover Principal scientist, National Institute for Plant Biotechnology, Lal Bahadur Shastri Building, Pusa Campus, New Delhi, India-110012 3. Mrs. Suchitra Pushkar Division of Plant Physiology, Indian Agricultural Research Institute, Pusa Campus, New Delhi, India-110012 4. Dr. Shashi Bala Singh, Division of Agricultural Chemistry, Indian Agricultural Research Institute, Pusa Campus, New Delhi, India-110012.

## Abstract

*Alternaria brassicae,* a necrotrophic pathogen causes Alternaria blight in members of the Brassicaceae family. An extensive yield-oriented breeding strategy has rendered Indian mustard (*Brassica juncea*) susceptible to many fungal pathogens however, Alternaria blight is one of the most pressing challenges of all because it causes almost 40-60% yield loss. Variable degree of tolerance is reported in wild relatives of *B. juncea, Sinapis alba* (White mustard) and *camelina sativa* (False flax) have been reported to exhibit moderate and high tolerance respectively against *A. brassicae*. Phytohormones are the essential regulator of the intricate mechanism of plant defence response. The hormones salicylic acid (SA) and jasmonic acid (JA) have been widely studied and recognized as important regulators of plant immune response. In the last decade, research has pointed out that other hormones like abscisic acid (ABA) also participate equally in plant defence. However, the role of ABA in defence responses and its cross-talk with SA and JA has not been fully understood in terms of *Brassica-A. brassicae* system. In this investigation, three genotypes-*B. juncea*, *S. alba* and *C. sativa* were selected and their response to exogenous application of SA, JA and ABA and their combination with *A. brassicae* were studied. Disease assessment, gene expression analysis and quantitative estimation of phytohormones showed that the *B. juncea* exhibited a weak JA-mediated defence response against *A. brassicae* and synergy between SA-ABA shifted the signalling mechanism to SA-mediated response leading to susceptibility in *B. juncea*. Tolerant genotypes, *S. alba* and *C. sativa* exhibited a robust JA-mediated response against *A. brassicae* and JA-ABA was found antagonistic in *Brassica-A. brassicae* phyto-pathosystem.

## 1. Introduction

Indian mustard (*Brassica juncea* L. Czern) is the third most important oilseed crop in the world and a leading oilseed crop of India, which contributes more than 80% to the total rapeseed-mustard production (Singh et al., 2017). Among all the members of the rapeseed-mustard group, *B. Juncea* alone is responsible to supply nearly 27% of vegetable oil to India and it is cultivated on approximately 7 million hectares of land (Rai et al.,2016). During its growth period, *B. juncea* is heavily challenged by numerous fungal pathogens and it is susceptible to many fungal pathogens including *Alternaria brassicae. A. brassicae* is a necrotrophic pathogen causing Alternaria blight in Brassica crops. The yield loss caused by this deadly pathogen has been estimated to be as high as 50 % depending on the severity of the disease incidence (Savary et al., 2006). The sessile nature of plants puts them in constant danger from thousands of pathogens but only a handful of them can colonise and spread disease. To fight immense pathogen pressure, plants are equipped with an intricate immune system inducing both constitutive (physical and chemical barriers) as well as induced immune responses (Kunkel and Brooks, 2002). The typical defence pathway activated by the plants in response to a pathogen attack depends on the lifestyle of the pathogen, i.e., whether the pathogen is a biotroph or a necrotroph. Biotrophic fungal pathogens derive their nutrients from live host cells and produce effectors to suppress the plant defence response (Derksen et al., 2013). On the other hand, necrotrophic pathogens kill the plant tissue with their weapons of toxins and chemicals before consuming them (Glazebrook, 2005). The perception of each type of pathogen and its molecular patterns determines the specific signal transduction route induced in the plants to tackle the enemy (Meng and Zhang, 2013). In response to a biotrophic pathogen, pathogen effectors are recognized by the resistance (R) genes in plants causing SA-induced signalling which leads to hypersensitivity response (HR) and cell death to check the further spread of the pathogen (Pieterse et al., 2012). Many studies have confirmed the SA-mediated defence response in the event of a biotrophic pathogen attack. For instance, mutants that fail to accumulate SA (such as *sid2*) or the SA-insensitive mutant *npr-1* exhibit higher susceptibility to biotrophs (An and Mou, 2011). This defence mechanism, however, will work in favour of necrotrophic pathogens and help them to colonize and spread therefore, plants evolved an alternate mechanism of resistance that is mediated by JA signalling which activates downstream defence responses caused by the MAPK-signalling cascade (Pieterse, 2008; Balbi and Devoto, 2008). Many reports demonstrated that mutation in JA receptor protein *COI 1* (Coronatine insensitive 1) gene renders the plants susceptible to necrotrophic pathogens like *A. brassicicola* and *B. cineria* (Thomma et al., 1999; Pandey et al., 2016). Therefore it is evident that initial recognition and signalling mediated by phytohormones during the early exposure to the pathogen is critical for the overall defence mechanism executed by the plants against the pathogen attack.

Phytohormones are the essential regulators of the defence responses in plants, The hormones SA and JA are recognized as the most important signalling molecules of the plant immune response. These two classes of hormones are believed to form the backbone of the plant immune responses (Derksen et al., 2013). The antagonistic nature of SA and JA pathways affecting the overall immune response in plants has been well studied and reported in many crop plants (Clarke et al., 2000; Penninckx et al., 1998; Flors et al., 2008). Research also shows that phytohormones which were typically known as growth hormones like cytokinin (CK) and auxin (aux) and abiotic stress hormone abscisic acid (ABA) also function in plant defence by influencing the SA-JA pathway (Shigenaga and Argueso, 2016). Therefore to understand the various factors influencing the overall immune response of plants it is vital to explore the influence of other phytohormones on the SA-JA interplay. Apart from its role in drought signalling in plants, ABA has emerged as an equally relevant player in signalling during biotic stress (Seilaniantz et al., 2011). The antagonism or synergism among other hormones and the SA-JA backbone contributes to the overall immune response of a plant against a specific pathogen. However, the role of ABA in cross-talk with SA-JA pathways is not clearly understood and there have been few reports suggesting contrasting outcomes. In biotrophic pathogens, it was reported that ABA treatment causes increased susceptibility in *Arabidopsis* (Ding et al., 2016) and rice (Xu et al., 2013). In terms of the necrotrophic pathogen, Anderson et al., (2004) reported an antagonistic interaction between ABA-JA causing increased susceptibility towards a necrotrophic fungal pathogen *Fusarium oxysporum* in *Arabidopsis.* A similar observation was reported in tomato where the exogenous application of ABA leads to increased susceptibility toward necrotrophic pathogen *Botrytis cinerea* (Audenaert et al., 2017). On the contrary, few researchers suggested that ABA treatment causes increased resistance toward necrotrophic pathogens in rice (de Vleesschauwer et al., 2010) and in *Arabidopsis* (Fan et al., 2009). However, it is clear from the above case studies that the ABA, SA and JA orchestrate a specific chain of signalling which is determined by various factors like pathogen type, virulence and other physiological factors. Therefore the interplay of these hormones is specific to each phyto-pathosystem and manifestation of the cross-talks among hormones is a crucial factor for plant defence response which can determine its overall response to the pathogen.

In the light of the above information, our study was designed to evaluate the effect of individual hormones (SA, JA and ABA), and its combined effect with *A. brassicae* infection on the defence responses executed by three different genotypes-*B. juncea* (susceptible), *S. alba* (moderately tolerant) and *C. sativa* (highly tolerant). The idea behind the study was to investigate the effect of complex treatments on hormone biosynthesis in a susceptible genotype and then cross-compare it with the responses generated in a tolerant genotype to identify key signalling details responsible for tolerance toward *A. brassicae*. Three selected genotypes, *B. juncea* (susceptible), *S. alba* (moderately tolerant) and *C. sativa* (Highly tolerant) were subjected to three sets of treatments-Hormones (SA, JA and ABA), *A. brassicae* (spore suspension) and Hormone + Infection (SA, JA and ABA treatment each followed by *A. brassicae* inoculation). Gene expression analysis targeting major biosynthesis genes of SA, JA and ABA biosynthesis pathways, as well as downstream responsive genes, was performed. To further confirm our findings, we performed quantitative profiling of each hormone in all the samples using HPLC and GC-MS. Disease establishment and lesion formation data were also collected and analysed for all the combination samples to provide a complete phenotypic and molecular analysis of the signalling pathways involved in disease signalling. To the best of our knowledge, this is the first report on the cross-talk of ABA with SA and JA in *Brassica-A. brassicae* phyto-pathosystem and its cross-comparison between the susceptible/resistant genotypes.

## 2. Material and methods

### 2.1 Plant material and experimental setup

Three closely related Brassicaceae genotypes namely, *B. juncea* (susceptible)*, S. alba* (moderately tolerant) *and C. sativa* (Highly tolerant) were selected for this study based on the previous study reporting their variable tolerance towards *A. brassicae*. (Sharma et al., 2002).

Plants of each species were grown in a controlled environment growth chamber programmed for 16h/8h of the light/dark cycle at a temperature of 24^0^C/20^0^C for day/night and relative humidity of 90%. 45 days old plants of each genotype were subjected to various treatments-hormone treatment (exogenous application of SA, JA and ABA), pathogen inoculation (*A. brassicae* spore suspension) and combination of hormone and pathogen inoculation (Hormone + infection). leaf tissue samples for RNA isolation and hormone quantification were harvested at 0, 3,6,12, 24, 48,72 and 96 hours post-treatment. A mock set was included simultaneously for each set of treatments and three replicates were maintained for each treatment.

### 2.2 Hormone solutions preparation and treatments

Working solutions of the three hormones were prepared in ddH_2_O. SA (0.5 mM), JA (100<u>μM</u>) and ABA (100<u>μM</u>) and were used for exogenous treatments. Plants were placed inside the glass chambers for the hormone treatments and sprayed with the hormone solutions using a fine nozzle spray bottle to achieve uniform distribution of the solution. One glass chamber was dedicated for the treatment of one hormone and a separate glass chamber was used for mock treatment. Chambers were sealed immediately after the treatment using polythene sheets and were only briefly opened for the sample collection. Leaf tissues were collected as samples at fixed time intervals, the leaf was cut at the joint of the petiole and wrapped in aluminum foil and snap-frozen using liquid nitrogen and stored in a deep freezer until further use. These samples were used for RNA extraction and hormone quantification.

### 2.3 Pathogen culture and inoculation

*A. brassicae* pure culture was sub-cultured on potato dextrose agar (PDA) at 25± 2°C. 15 days old culture showing profuse sporulation was used for inoculation. The plate was flooded with distilled water and spores were collected by mildly scraping the colony surface. The suspension was filtered with two layers of muslin cloth and the filtrate was adjusted to a spore concentration of 5×10^5^ ml^−1^ with the help of a hemocytometer. A surfactant, tween 20 (0.05%) was also added to the suspension to maintain the surface tension on the leaf surface which helps the spore stick to the leaf. Inoculation of the plant was done by following the drop inoculation method (Giri et al., 2014). For the hormone + Infection experiment, plants were sprayed with the respective hormones and the pathogen inoculation was subjected after 1hr of the hormone spray and the samples were collected at a fixed time interval thereafter. Control plants treated with DDW were harvested simultaneously for all the treatments.

### 2.4 Disease progression analysis

To understand the effect of phytohormones (SA, JA and ABA) on disease establishment and progression over time a polyhouse experiment was conducted for all three-host species (*B. juncea, S. alba* and *C. sativa*). Plants of each genotype were planted in large pots of diameter 20” in a polyhouse and were subjected to the following treatments: Mock (ddH_2_O), Inoculation (*A. brassicae* spore suspension), SA+I (exogenous SA treatment followed by inoculation), JA+I (exogenous JA treatment followed by inoculation) and ABA+I (exogenous ABA treatment followed by inoculation). For each treatment, five replicates were maintained and the disease progression was monitored at an interval of 5 days after inoculation until 30 days. Observations like days to 1^st^ symptom appearance, number of leaves with lesion per plant and lesion diameter were recorded. Polyhouse conditions were kept highly favourable for disease development as established by (Meena et al., 2004) - 18-27°C of maximum temperature, 8-12°C of minimum temperature, >90 % relative humidity (RH) during morning and nights, >70% RH during afternoons and >10h of leaf wetness in the previous week was maintained using overhead sprinklers. The average data of the above parameters were used to understand the disease progression trends in all three genotypes.

### 2.5 RNA isolation, cDNA synthesis and quantitative real-time PCR analysis

The TRIzol^TM^reagent (Invitrogen) was used for total RNA extraction from frozen leaf tissues. The purity and quantity of RNA were determined using the NanoDrop^TM^2000 spectrophotometer (Thermo scientific). First-strand cDNA synthesis kit (Thermo Scientific) was used to synthesize the first-strand cDNA from 1µg of RNA for 20 µl of reaction with oligo dT primers. cDNA was diluted to 1:10 (cDNA: nuclease-free water) for further qPCR reactions. To design the primers of genes, the *Arabidopsis* CDS sequence was obtained from Genbank and used as a query sequence in BlASTn to obtain the homologous sequences from *B. juncea* and its wild relatives from the B. *rapa* genome portal (http://brassicadb.org/brad).

To ensure positive amplification in *B. juncea* and both the wild relatives, sequences of each gene were aligned using the clustalw tool and primers were designed from the consensus region of the aligned sequences. For all the genes, primers were designed using the online tool Primer 3 (Untergasser et al., 2012) with the following parameters: primer length 20-25 bp, amplicon size 100-200bp, GC content 60-65%, melting temperature 60-65^0^C, absence of hairpin structure, homodimer and heterodimer.

Details of primer sequences used for qRT-PCR analysis are provided in Table 1. qRT-PCR was performed using SYBR green technology in 96 well optical plates with lightcycler 480 real-time PCR machine (Roche) adopting the following thermocycle conditions: 95^0^C for 5min, 40 cycles of 95^0^C for 20 sec, 60^0^C for 60 sec and 72^0^C for 30 sec. PCR reaction mixture was prepared at a final volume of 20µl which contains 10 µl of TB Green Premix Ex Taq II (Takara), 10pM of forward and reverse primers and 2µl of diluted cDNA. NTC (no template control) was maintained in every run for all the reactions. The most accurate reference gene was selected based on the previous reports on normalization studies on the selected genotypes (Dixit et al., 2019, 2020a)

**Table 1.**
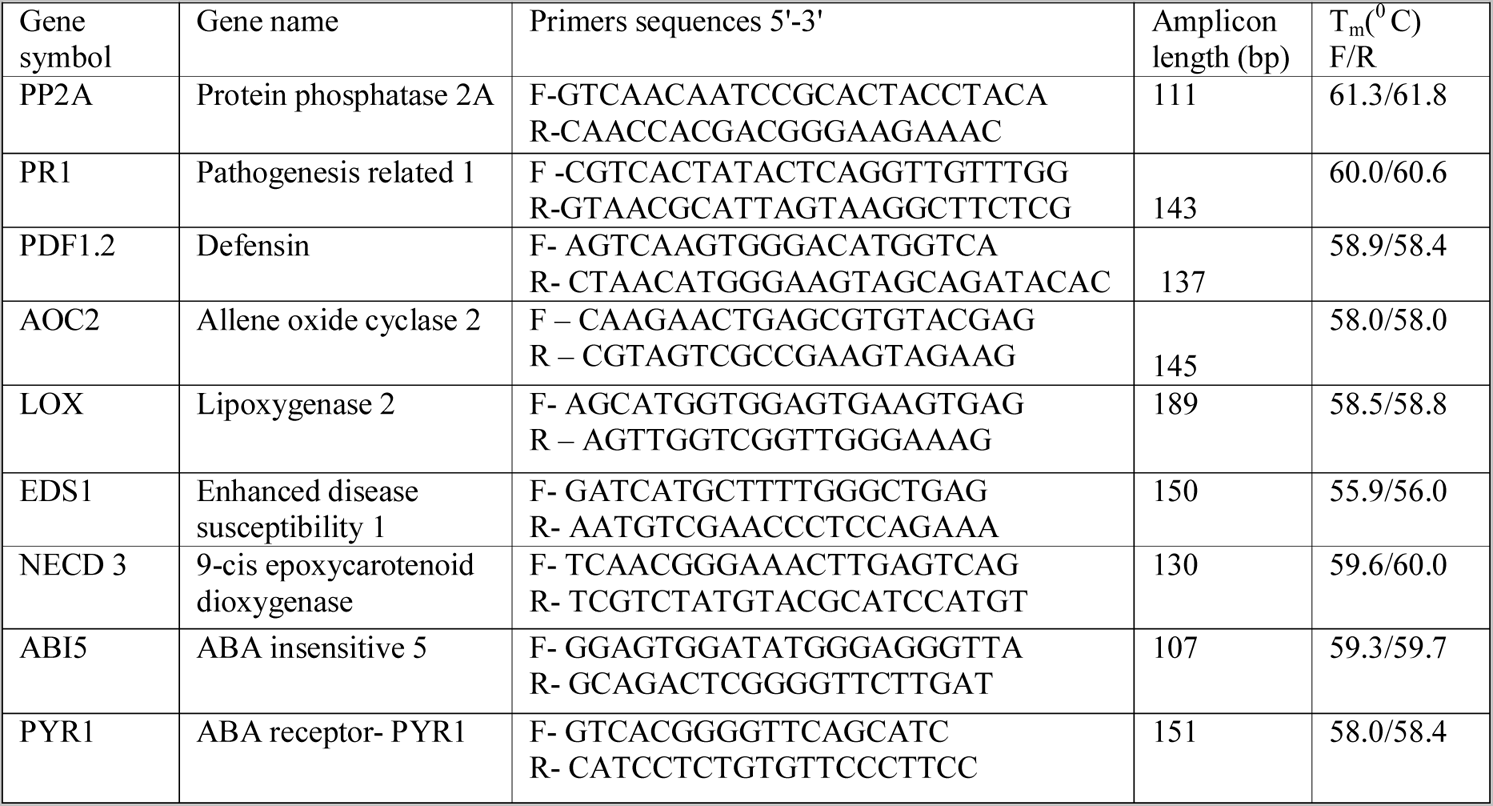
Details of the primers used in the qRT-PCR analysis.

### 2.6 Phytohormones extractions and quantifications using HPLC and GC-MS

Leaf samples were used for hormone extraction. SA and ABA were extracted using a solvent-based extraction procedure and the quantification was done using HPLC. JA being a volatile phytohormone was extracted using a solid-phase extraction procedure involving SPE cartridge and quantification was done using GC-MS. The extraction procedure followed for each hormone and its detection conditions is s mentioned below:

**2.5.1 ABA extraction and quantification**: Abscisic acid extraction protocol was derived from the protocol mentioned in (Nakurte et al., 2012). Briefly, 1g of leaf tissue was crushed in Liq N_2_, pulverized and stored at −80^0^C. Frozen samples were homogenized with 10 ml of extraction buffer (80ml of acetone, 1 ml of glacial acetic acid and 100mg of 2,6 di-tret-butyl 4-methyl phenol in a total volume of 100ml). The extraction was repeated thrice with 10 ml extraction buffer by centrifugation @ 9800 g for 10 min each time. The homogenate was filtered using Whatman No 1 filter paper and the solvent was evaporated with the help of a boiling rotary evaporator. As the acetone evaporated, the lipid-soluble component of the samples was deposited on the walls of the boiling flask. The precipitate was dissolved in 1% acetic acid solution to give an amber-colored aqueous solution which was used for quantification. HPLC conditions and detector were kept as follows: variable wavelength detector wavelength was set at 265 nm. Zorbax Eclipse XDB-C18 column from Agilent technologies of dimension 4.6 × 250mm, 5μ pore size and temperature of the column was maintained at 30^0^C. The mobile phase used for the analysis was 1% acetic acid in 95% of methanol at a flow rate of 1ml/ min. The injection volume was set at 20 μl.

**2.5.2 SA extraction and quantification**: Salicylic acid extraction protocol was derived from the protocol used by (Li et al., 1999). 1g of frozen leaf tissue was crushed in Liq N_2_ and homogenized with 3ml of 90% methanol, homogenized samples were sonicated in a sonicator bath for 20 min and then centrifuged @ 7500g for 20 min. The supernatant was collected in a fresh tube and the leaf pellet was re-extracted with 100% methanol following the same steps as above. Supernatants collected from both the steps were mixed thoroughly and divided into two equal parts. One half was used to extract conjugated SA and the second half was used for the extraction of unconjugated SA. For unconjugated SA extraction 3ml of 5% Trichloroacetic acid (TCA) was added to almost dry samples and subjected to a vigorous vertex. For conjugated SA extraction, the sample supernatant fraction is subjected to acid hydrolysis by 1ml of 2M HCl for 60 min @ 65^0^C. Both the fractions were then extracted twice with an equal volume of extraction buffer (ethylacetate: cyclopentane: isopropanol - 100:99:1) further the dried samples were dissolved in methanol. Samples were filtered and run using 90% methanol as the mobile phase and A florescence detector was used for the analysis with an excitation/ emission wavelength of 295/405 nm. Zorbax Eclipse XDB-C18 column from Agilent technologies of dimension 4.6 × 250mm, 5μ pore size was used and temperature of the column was maintained at 35^0^C. A flow rate of 1ml/ min. The injection volume was set at 20 μl.

**2.5.3 JA extraction and quantification:** Direct Solid-phase extraction column method modified by (Liu et al., 2010) was used for the extraction of Jasmonic acid. Briefly, 1g of frozen leaf tissue was crushed in Liq N_2_ and then homogenized using 5 ml of 100% methanol and kept at 4^0^C overnight. Samples were centrifuged @ 4800g for 10 min and the supernatant was transferred to a fresh tube. The leaf pellet was re-extracted by 3ml of methanol and both the supernatant were combined. 3ml of HPLC grade H_2_O was added to the combined mixture.

The solution was applied to the Waters Sep-pak C18 cartridge. The cartridge was washed with 2 ml of 20% methanol and 1 ml of 30% methanol and finally eluted with 1 ml of 100% methanol. The eluted solution was collected as the samples extract ready for analysis. GC-MS conditions were standardized and applied based on the conditions listed in (Meyer et al., 2003). Sampled volatiles was analyzed by GC on a Shimadzu model 17A instrument equipped with an auto-injector and an FID. Capillary columns with (30 mm× 0.25 mm and 0.25 mm film thickness) were used. Helium was used as the mobile phase and the peaks of all the prospective compounds were manually integrated and peak area was calculated. Retention time for the JA was estimated based on the peak obtained by the standard solution. The registered peak was run through NIST and wiley libraries for accurate analysis. The actual quantification was performed using standard peak area obtained by the standard curve generated by known concentration of the standard solution.

## 4. Results

### 4.1 The effect of hormone treatments on disease establishment and progression

As reported earlier*, B. juncea* is susceptible to the pathogen *A. brassicae* (Sharma *et al*., 2002). In our investigation, we observed an early establishment of disease *in B. juncea* as compared to the tolerant wild genotypes. In *B. juncea* pin-headed lesions can be observed as early as 24 hrs after inoculation (hai) and the lesions grew rapidly and starts to spread within 96 hai. The exogenous application of hormones SA, JA and ABA showed a clear effect on the overall disease progression. As evident from figure1A, ABA and SA treatment, in *B. juncea* significantly aggravated the disease symptoms resulting in higher tissue damage and increased disease spread. As compared to *A. brassicae* inoculated samples the lesion size and surrounding tissue chlorosis were higher in ABA and SA pre-treated plants followed by A. brassicae inoculation. On the other hand, JA pre-treated samples showed reduced lesion size and restricted spread after inoculation. As shown in figure 1B, the average lesion size observed in *A. brassicae* inoculated samples 96 hai was 0.76 cm whereas, the average lesion size in ABA and SA treated samples were 1.72 cm and 2.2 cm respectively. The vigorous spread of lesions in ABA and SA treated samples indicates that these hormones aided in the spread and establishment of the pathogen and intensified disease symptoms in *B. juncea*. Contrastingly, JA pre-treated plants managed to check the spread of the lesions and developed small-sized lesions with lesser damage to the surrounding leaf tissues (figure1A).

**Figure 1:**
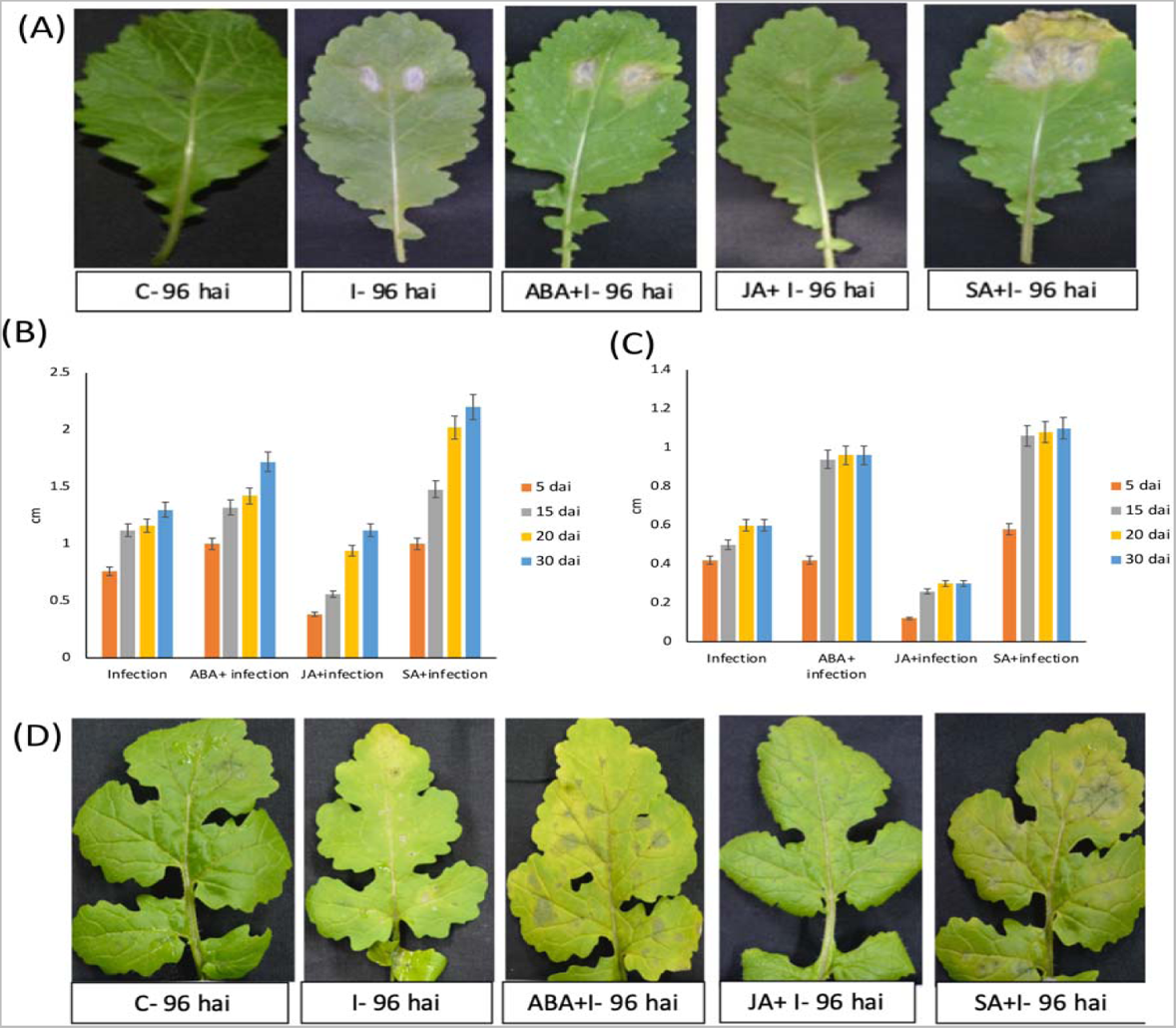
Effect of hormone treatment on disease establishment and progression in *B. juncea* (susceptible), and *S. alba* (moderately tolerant). (A) Effect of ABA, JA and SA pre-treatment on lesion size and leaf tissue damage in *B. juncea* 96 hai with *A. brassicae.* (B) Graph showing the disease progression trend observed in ABA, JA and SA pre-treated *B. juncea* plants from 5 days after inoculation (dai) to 30 dai as compared to the inoculated plants (without hormone pre-treatment) (C) Graph showing the disease progression trend observed in ABA, JA and SA pre-treated *S. alba* plants from 5 dai to 30 dai as compared to the inoculated plants (without hormone pre-treatment). (D) Effect of ABA, JA and SA pre-treatment on lesion size and leaf tissue damage in *S. alba* 96 hai with *A. brassicae*.

In general, the tolerant genotypes *S. alba* and *C. sativa* exhibited less aggravated disease symptoms as compared to the susceptible genotype *B. juncea.* Although, the exogenous hormone treatment did influence the overall disease incidence in tolerant genotypes the effect was less dramatic (figure1D and 2A). As shown in figure 1C and figure 2B, the average lesion size in *S. alba* and *C. sativa* was 0.42 cm and 0.26 cm respectively 96 hai which is significantly lower than the average lesion size in *B. juncea* (1.12cm). Upon exogenous hormone treatment, it was observed that ABA and SA treatment aggravated the pathogen spread but the tolerant genotypes manage to keep the damage to the minimum and quickly suppressed the further spread of the disease as evidenced by the trend observed in average lesion size in the later stages of our experiment. The average lesion size increase was insignificant from 15 dai to 30 dai (Figure 1C and 2 D) in both ABA and SA treated plants. This indicated that the immediate effect of disbalance in the phytohormone system of tolerant genotypes worked in the favour of the pathogen. However, the effect was quickly nullified and the further spread of the pathogen was restricted. This phenomenon was found weak in susceptible *B. juncea*. Interestingly, in tolerant genotypes, JA treatment greatly reduced the disease symptoms and a significant delay in disease establishment was noticed in tolerant genotypes as compared to the susceptible genotype (Figure 1C and D and Figure 2). These results indicated a strong JA mediated response against *A. brassicae* exhibited by the tolerant genotypes could be the reason for delayed infection and lower disease spread and *B. juncea* exhibited a weaker JA response towards the pathogen.

**Figure 2:**
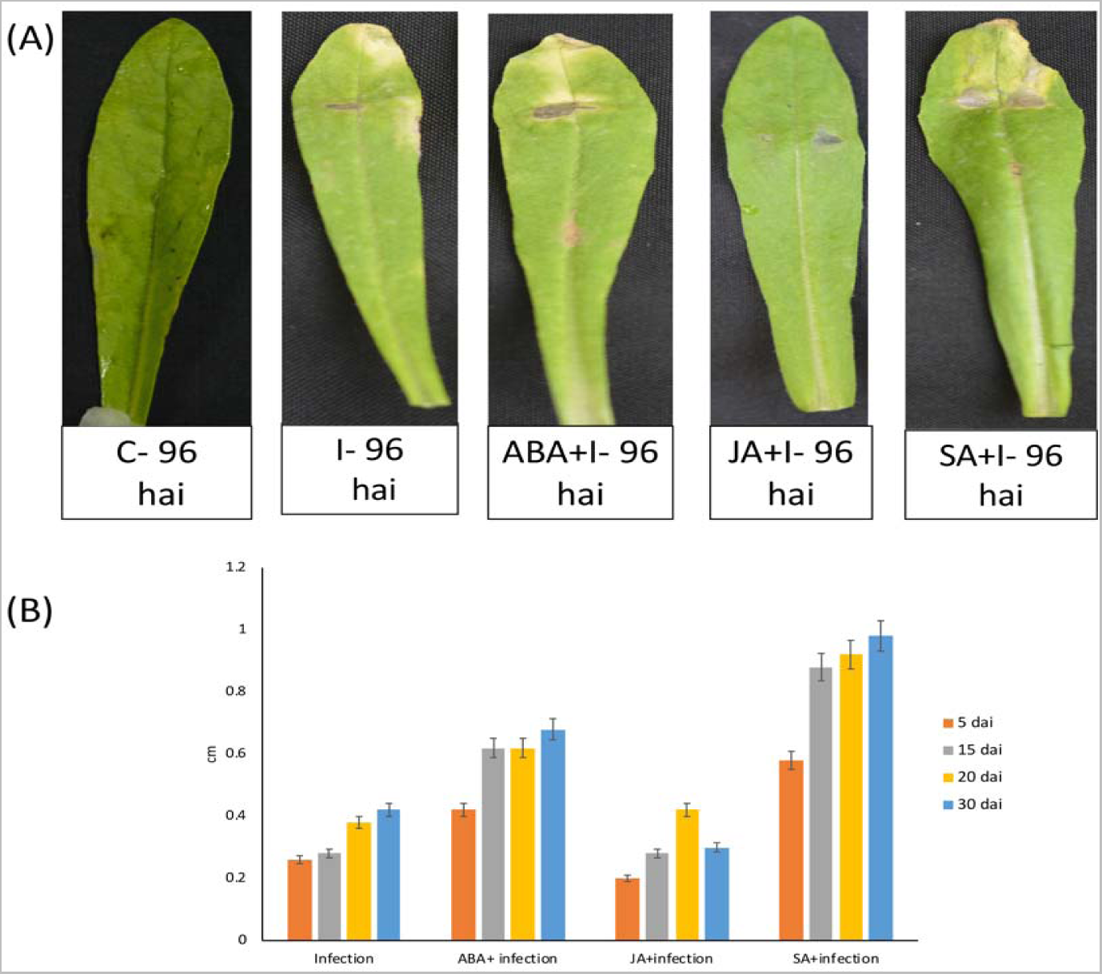
Effect of hormone pre-treatment (ABA, JA and SA) on disease establishment and progression in highly tolerant genotype *C. sativa*. (A) The difference in lesion size and leaf tissue damage shown in hormone pre-treated plants of highly tolerant genotype *C. sativa* 96 hai with *A. brassicae*. (B) Graph showing the disease spread in tolerant genotype *C. sativa* after hormone (ABA, JA and SA) pre-treatment plant as compared to inoculated plants (without hormone pre-treatment) from 5 dai till 30 dai.

### 4.2 Gene expression analysis of ABA, JA and SA biosynthesis genes, defence responsive genes and hormone quantification analysis

As evident from the disease progression analysis it is clear that JA pre-treated plants exhibited a strong defence response against *A. brassicae* in all three genotypes. However, in susceptible *B. juncea,* the tolerance response quickly succumbed and the overall effect on the disease intensity remained unaffected at the later stages of the infection. This indicated that the *B. juncea* is unable to maintain a higher cellular concentration of JA to support the threshold defence response whereas, tolerant genotypes maintain a higher cellular concentration of JA enough to trigger a strong defence response. It was also clear from the disease progression analysis that SA and ABA are working in favour of the pathogen and causing increased disease incidence and to confirm these results we performed a comparative analysis of the expression pattern of hormone biosynthesis genes of three hormones under investigation -ABA, JA and SA in response to various treatments among *B. juncea*, *S. alba* and *C. sativa.* We shall discuss our findings concerning each hormone signalling pathway and its effect on disease progression in the following sections:

#### 4.2.1 Expression profile of JA biosynthesis pathway genes

The plant hormone jasmonate (JA) regulates diverse aspects of plant growth, development, and immunity. This hormone plays a critical role in controlling defence responses to an extraordinary range of biotic aggressors, most notably necrotrophic pathogens (Peña-Cortés et al., 2004). JA synthesis is initiated in the chloroplast, where linolenic acid is converted to 12-oxo-phytodienoic acid (OPDA) by the sequential action of lipoxygenase (LOX), allene oxide synthase (AOS), and allene oxide cyclase (AOC). After pathogen inoculation, the up-regulation of the JA-responsive genes such as plant defensin 1.2 (*PDF1.2*) and vegetative storage protein 2 (*VSP2*), provides evidence that JA acts as a signalling molecule in plant immunity to necrotrophic pathogens (Balbi and Devoto, 2008). We analysed the expression of two major biosynthesis genes of JA, *LOX2* and *AOC2* selected based on the previous reports which suggested that these two genes are the major JA involved in defence signalling responses (Spoel et al., 2003a, Wasternack and Song, 2017)

It was observed that SA and ABA treatment downregulated the *LOX2* gene expression in all three genotypes. This clarifies that SA and ABA act as negative regulators for JA biosynthesis genes (Figure 3). *A. brassicae* inoculation did not affect the expression of the *LOX2* gene in *B. juncea* however, in *S. alba* and *C. sativa*, *LOX2* gene expression raised to approximately 2-fold 12 hrs after inoculation and the higher expression level was maintained till 24 hrs after inoculation. MeJA treated samples exhibited a 6-fold increase in expression 12 hrs after treatment in *B. juncea* and in tolerant genotypes the expression level increased to almost 10 folds. Interestingly in the hormone + infection sample set, ABA and SA pre-treated samples continue to show downregulation of the *LOX2* gene even after inoculation in all three genotypes. On the other hand, the MeJA pre-treated samples showed a 3-fold increase in *LOX2* gene expression 12 hrs after inoculation in *B. juncea* and the level of expression in tolerant genotypes was found to be approximately 4-fold. These observations indicated that *S. alba* and *C. sativa* exhibited a quick induction in JA biosynthesis in response to *A. brassicae*. Whereas, *B. juncea* failed to execute an efficient signalling strategy in response to *A. brassicae*.

**Figure 3:**
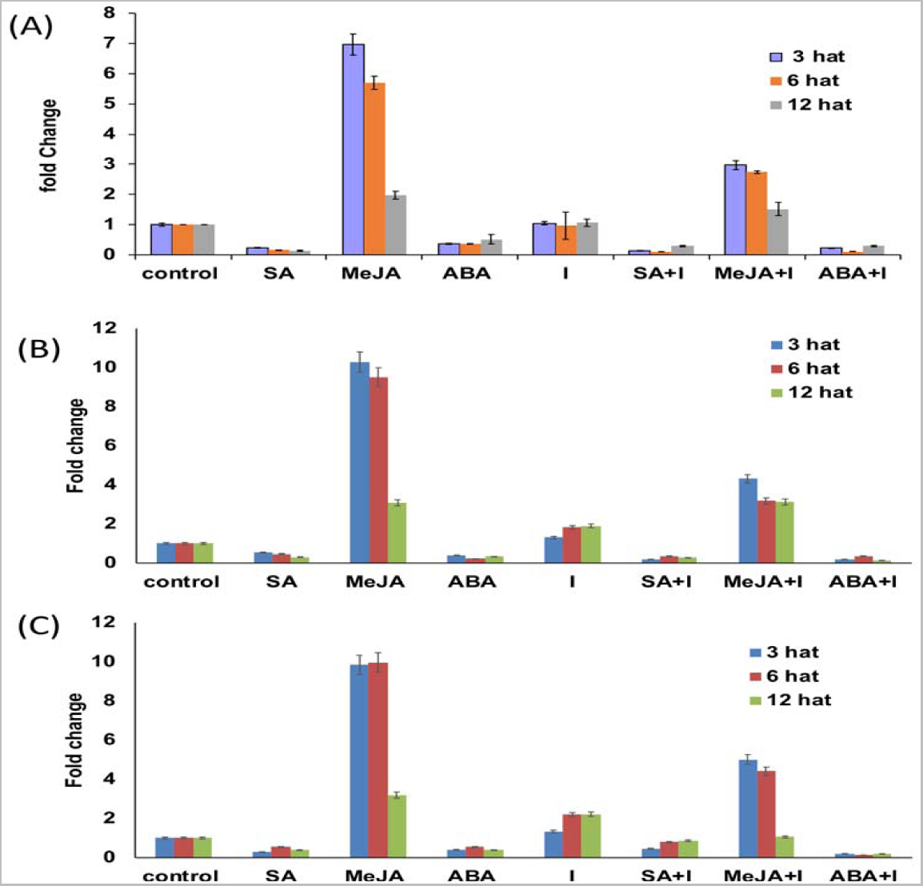
Expression profile of *LOX2* gene in all three genotypes. Transcript levels of *LOX2* gene in response to various treatments at 3, 6 and 12 hrs after treatment (hat) in (A) *B. juncea* (B) *S. alba* (C) *C. sativa.* In this figure the treatments are- salicylic acid (SA), methyl jasmonate (MeJA), abscisic acid (ABA), I (Infection, *A. brassicae* spore suspension was used in this treatment), SA+I (SA pre-treatment followed by *A. brassicae* inoculation), MeJA+I (MeJA pre-treatment followed by *A. brassicae* inoculation) and ABA+I (ABA pre-treatment followed by *A. brassicae* inoculation).

In the *AOC2* gene, we observed similar effects of hormone treatment and *A. brassicae* inoculation on the expression pattern as observed in *LOX2* with few exceptions. In *B. juncea*, a 1.6-fold induction was observed in *A. brassicae* inoculated samples 3 hr after inoculation no such induction was observed in LOX2 gene expression in *B. juncea* (Figure 4A). This indicates that *AOC2* is linked closely to the signalling pathway induced in response to *A. brassicae*. In tolerant genotypes, 4-fold induction was observed in *AOC2* gene expression in response to the *A. brassicae* inoculation (Figures 4 B and C). The early and efficient induction of *AOC2* genes in tolerant genotypes in response to *A. brassicae* further confirmed that among all the JA biosynthesis genes, the *AOC2* gene is closely connected to defence signalling network. Similar to *LOX2* gene expression, SA and ABA acted as negative regulators of *AOC2* gene expression. This confirmed that ABA and JA are antagonistic in *A. brassicae*-*Brassica* phytopatho system. It can also be understood from the comparative gene expression analysis of JA biosynthesis genes that tolerant genotypes have an efficient signalling network which quickly upregulates the biosynthesis of JA and triggers a JA mediated defence response on occasion of *A. brassicae* infection. On the other hand, in susceptible genotype *B. juncea,* this intricate signalling system is rendered weak.

**Figure 4:**
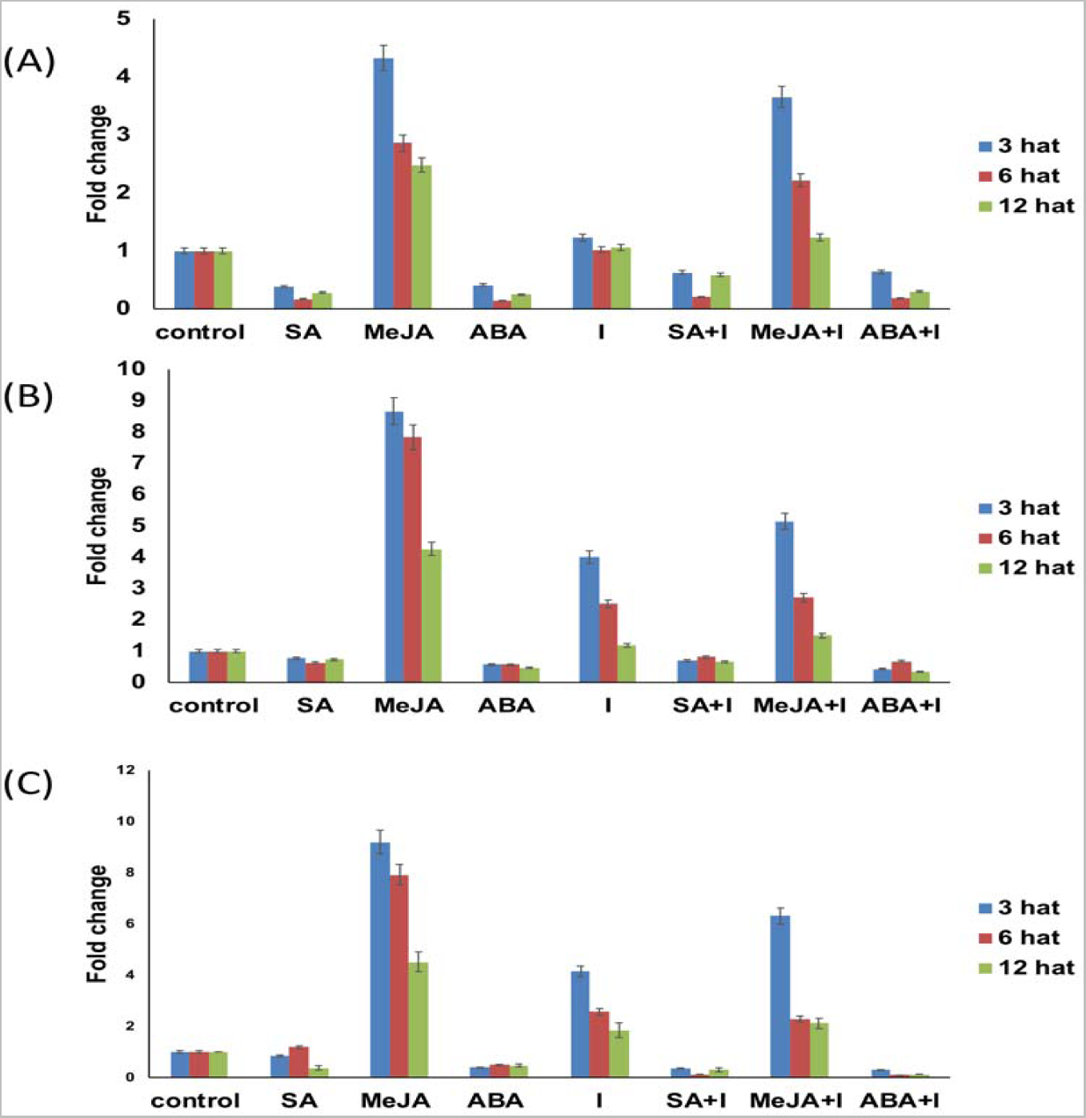
Expression profile of *AOC2* gene in all three genotypes. Transcript levels of *AOC2* gene in response to various treatments at 3, 6 and 12 hrs after treatment in(A) *B. juncea* (B) *S. alba* (C) *C. sativa.* In this figure the treatments are- salicylic acid (SA), methyl jasmonate (MeJA), abscisic acid (ABA), I (Infection, *A. brassicae* spore suspension was used in this treatment), SA+I (SA pre-treatment followed by *A. brassicae* inoculation), MeJA+I (MeJA pre-treatment followed by *A. brassicae* inoculation) and ABA+I (ABA pre-treatment followed by *A. brassicae* inoculation).

#### 4.2.2 Comparative gene expression analysis of PDF1.2 gene

The defence response in plants is executed by the activation of downstream responsive genes of signalling pathways. *PDF1.2* is an important JA responsive anti-fungal defensin gene and it confers tolerance toward necrotrophic pathogens in plants (Rowe et al., 2010). In our investigation, the MeJA and MeJA + infection treatment upregulated the expression of the *PDF1.2* gene in all three genotypes. However, the tolerant genotypes exhibited higher induction as compared to *B. juncea.* As shown in figure 5, MeJA treatment causes 7-fold induction in *PDF1.2* expression in *B. juncea* 3 hr after treatment whereas *S. alba* and *C. sativa* exhibited 8-fold and 10-fold induction respectively. In response to *A. brassicae* inoculation, *B. juncea* showed an upregulation of 2-fold at 24 hrs after inoculation (Figure 5A) whereas, both the tolerant genotypes showed an early and stronger induction of *PDF1.2* wherein, *S. alba* and *C. sativa* showed an upregulation of 5-fold and 7-fold respectively in the PDF1.2 transcript levels at 12 hrs after inoculation and maintained a high level of expression till 24 hrs after inoculation (Figure 5B and C). SA and ABA treated samples showed downregulation in *PDF1.2* gene expression in all three genotypes, confirming the antagonistic role of SA and ABA in JA-induced defense responses. Interestingly, In MeJA + I sample of *B. juncea* the expression level of *PDF1.2* was much higher at 3 hrs and 6hrs after inoculation and upon correlation with the disease establishment data we can say that due to exogenous MeJA application the cellular JA levels were higher at the early stages of the infection in these samples which makes it difficult for the pathogen to develop prominent symptoms the disease (Figure1A). Therefore, we can conclude that induction of JA synthesis in the early stages of the infection is critical for the pathogen infestation. The tolerant genotypes exhibited a quick surge in JA biosynthesis by causing an early induction of JA biosynthesis genes leading to a stronger defense response at early infection stages.

**Figure 5:**
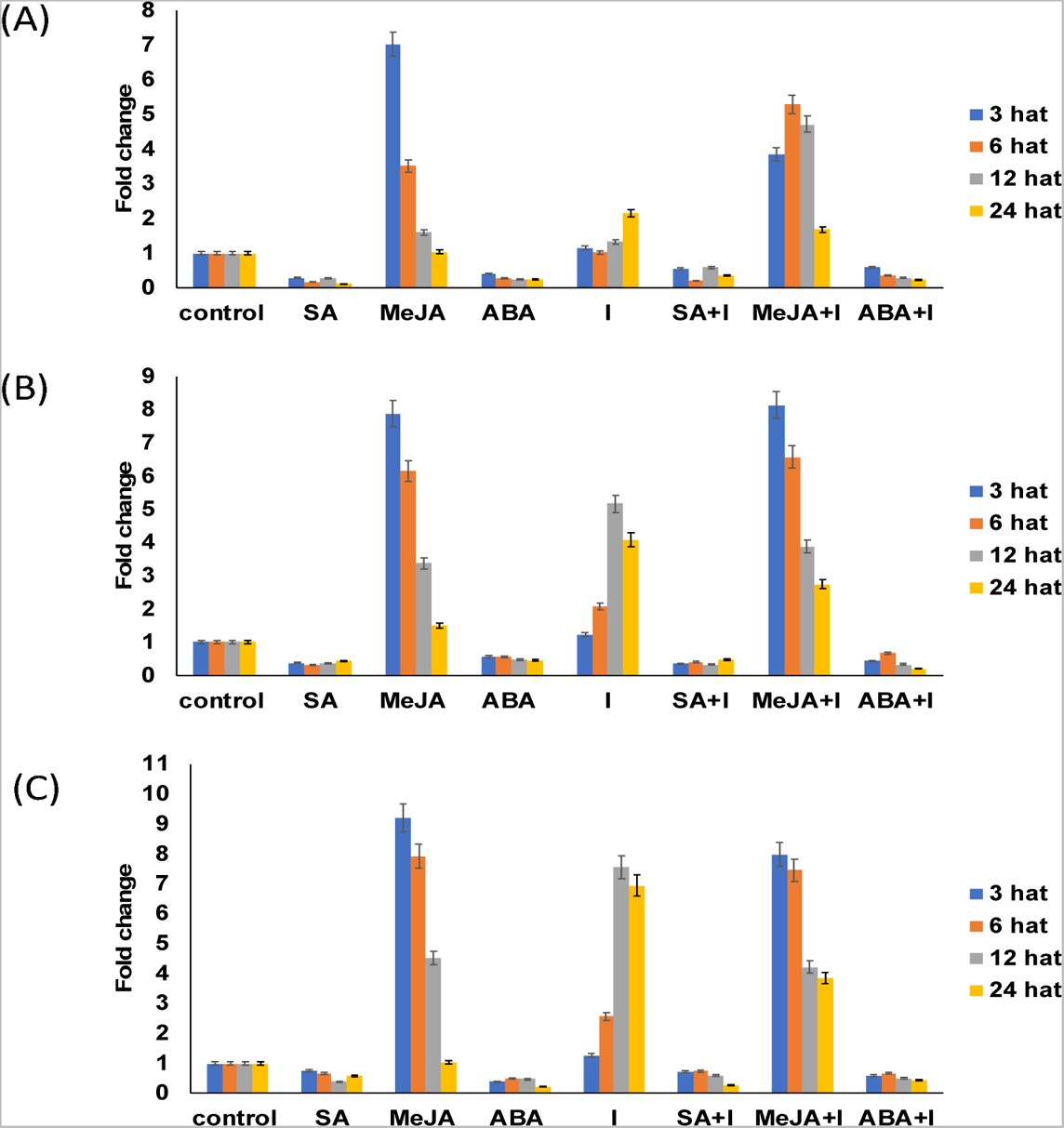
Expression profile of *PDF1.2* in all three genotypes. Transcript levels of *PDF1.2* gene in response to various treatments at 3, 6,12 and 24 hrs after treatment (hat) in (A) *B. juncea* (B) *S. alba* (C) *C. sativa.* In this figure the treatments are- salicylic acid (SA), methyl jasmonate (MeJA), abscisic acid (ABA), I (Infection, *A. brassicae* spore suspension was used in this treatment), SA+I (SA pre-treatment followed by *A. brassicae* inoculation), MeJA+I (MeJA pre-treatment followed by *A. brassicae* inoculation) and ABA+I (ABA pre-treatment followed by *A. brassicae* inoculation).

#### 4.2.3 JA quantification

Based on the gene expression profile of JA biosynthesis genes it is clear that tolerant genotypes were able to quickly deploy JA biosynthesis genes leading to early induction of JA synthesis which in turn led to efficient JA mediated defence response and this intricate signalling network between JA biosynthesis and defence signalling is found weak in susceptible genotype. To further confirm our results, we performed a quantitative analysis of JA concentration in all the sample sets of *B. juncea*, *S. alba* and *C. sativa* at different time intervals. The comparison of the relative JA content in susceptible/ tolerant genotypes under different treatments revealed the insights of JA, SA and ABA-mediated interplay. In SA treated samples, the JA concentration in susceptible and in tolerant genotypes was found lower as compared to the mock samples. However, in *B. juncea* the JA content dropped to almost 50% 3 hrs after treatment and could not recover till 24 hrs after treatment (Figure 6A). Whereas in *S. alba* and *C. sativa,* the drop in JA concentration was noticed but the recovery to the natural JA concentration was rapid (Figure 6A). ABA treatment drastically reduced the cellular JA concentration in all three genotypes however, *B. juncea* exhibited a maximum of 60% drop in JA level at 12 hrs after treatment. *C. sativa* maintained a fairly stable JA cellular level in response to ABA at all time points (Figure 6B). Interestingly, *A. brassicae* inoculated samples exhibited a dramatic JA profile where *B. juncea* showed lower JA levels after early hrs of treatment (3 and 6 hat) however a small surge in JA concentration (6μg/g leaf tissue) was observed between 12 and 24 hrs after inoculation (Figure 6C). Tolerant genotypes showed a significant increase in JA levels in response to *A. brassicae* inoculation at all time points (Figure 6C). The ABA+ Infection treatment caused a significant reduction in JA levels in susceptible *B. juncea* (Figure 6D). On the other hand, in tolerant genotypes, a reduction in JA level was noticed at 3hrs which rapidly normalized at the later time points. Based on the collective understanding from the disease progression analysis, gene expression analysis of JA signalling pathway genes and quantitative JA analysis it can be concluded that in response to *A. brassicae* the susceptible *B. juncea* exhibit slow signal perception and transmission leading to weak induction of JA biosynthesis and therefore exhibit a weaker JA mediated defence response against the pathogen. On the other hand, tolerant genotypes have evolved for early defence response against *A. brassicae*. They rapidly generate strong signalling streams to induce strong JA biosynthesis induction leading to a robust JA mediated defence response. From our findings, we confirmed that ABA-JA exhibit an antagonistic relationship in *Brassica-A. brassicae* phyto-pathosystem.

**Figure 6.**
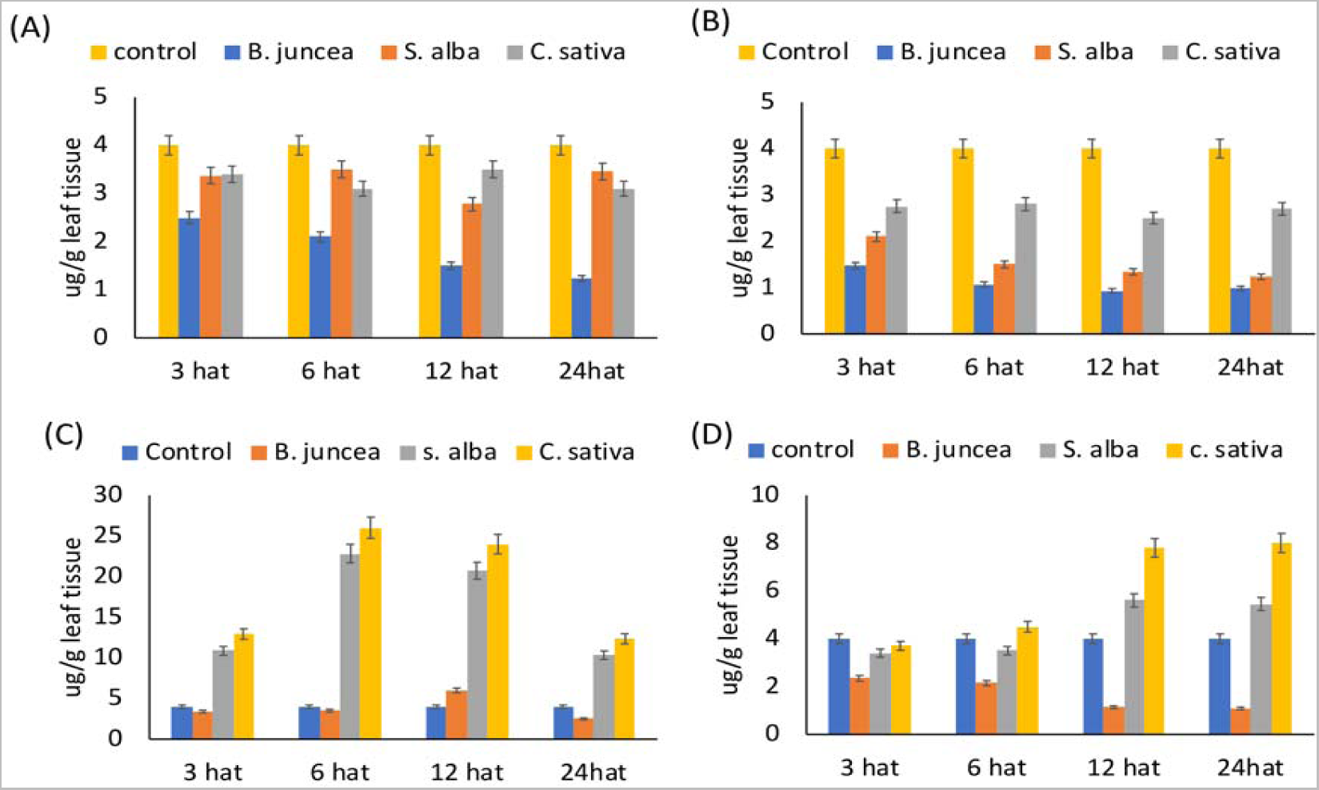
JA levels estimated in leaf samples of all three genotypes subjected to various treatments using GC-MS analysis. **(A)** Relative JA concentration estimated in per g leaf tissue of all three genotypes from 3 hat to 24 hat subjected to (A) SA treatments. (B) ABA treatments (C) *A. brassicae* inoculation. (D) ABA+I (ABA pre-treatment followed by *A. brassicae* inoculation). JA content in samples were calculated based on the standard curve generated from the area under peak obtained from the gradient concentrations of standard solution.

#### 4.2.4 Expression profile of ABA biosynthesis genes

ABA biosynthesis occurs mostly in plastids and is initiated by the conversion of zeaxanthin to trans-violaxanthin, by the enzyme zeaxanthin epoxidase (ZEP). This step is then followed by cleavage into xanthoxin by 9-cis-epoxy carotenoid dioxygenases (NCED), in what is considered the rate-limiting step of ABA biosynthesis. In the cytosol, xanthoxin is converted to abscisic aldehyde by a short-chain dehydrogenase/reductase (SDR) and then oxidized to ABA by the enzyme aldehyde oxidase (AAO). ABI5 is the transcription factor involved in ABA signaling in plants towards abiotic and biotic stresses. In this study, we analyzed the expression pattern of *NCED*3, *PYR1* and *ABI5* genes based on the previous reports suggesting the vital involvement of these genes in ABA-mediated signaling in response to biotic stresses (Fan et al., 2009, Xu et al., 2013).

The expression profile of *NCED3,* an important ABA biosynthesis gene was studied in different sample sets of all three genotypes as described in the previous section. MeJA and ABA treatments significantly downregulated the *NCED3* gene expression in all three genotypes. The downregulation of ABA biosynthesis genes after MeJA confirms the two-way antagonistic relation between JA-ABA in these genotypes. The end product of the ABA biosynthesis reduced the expression of ABA biosynthesis genes because many biosynthesis pathways are self-regulatory some reports suggested that cellular ABA concentration negatively regulates ABA accumulation (Qin and Zeevaart, 2002). SA treated samples upregulated *NCED3* gene expression up to 2-fold in all three genotypes and no significant variation in gene expression was observed among the susceptible and the tolerant genotypes.

However, based on this result it can be speculated that SA act as a positive regulator of the ABA biosynthesis in these *Brassica* genotypes. In the *A. brassicae* inoculated samples, *NCED3* expression was found upregulated to 4-folds at 6 hrs after inoculation and maintained a higher level of expression till 12 hrs after inoculation in both *B. juncea* (Susceptible) and *S. alba* (moderately tolerant). Interestingly, in highly tolerant genotype *C. sativa* only minor induction in *NCED3* expression, close to 1.8-fold was noticed 6 hrs after inoculation. These results can be explained based on the antagonist interaction between JA and ABA, as it was found by the JA quantification analysis that *C. sativa* exhibits higher cellular JA levels as compared to susceptible *B. juncea,* this could lead to suppression of *NCED3* gene expression and in turn negatively regulate the ABA production. The expression pattern of the *NCED3* gene in the hormone + infection sample set further gave us interesting insights into cross-talk between SA, JA and ABA. As expected the MeJA pre-treated samples of all three genotypes showed a consistent downregulation in *NCED3* expression in all three genotypes. SA and ABA pre-treated plants of *B. juncea* showed approximately 2-fold induction at 6 hrs after inoculation till 12 hrs after inoculation. Contrastingly in tolerant genotypes, SA pre-treated samples showed limited induction in *NCED3* expression and in ABA pre-treated samples a downregulation of gene expression was observed. These results can be explained based on the cross-talk dynamics between SA-ABA, SA pre-treatment causes the upregulation in *NCED3* expression after the inoculation process the cellular physiology of tolerant genotype shifts towards the strong JA mediated response and the sudden rise in JA levels after inoculation (as observed in the JA quantification analysis) leads to suppression of *NCED3* expression. However, the susceptible genotype fails to execute a strong JA signalling which shifts the cross-talk balance more towards an ABA-SA-based response which in turn work in the favour of the pathogen. These results can be further confirmed by the ABA quantification in all the sample sets.

No significant variation in the gene expression pattern of *ABI5* and *PYR1* genes were noticed among the susceptible and the tolerant genotypes in response to various treatments. However, there were a few exceptions susceptible genotype *B. juncea*, at 24 hr after inoculation approx. 1.5-2-fold change in ABI5 gene expression was recorded. Downregulation of ABI5 is noticed in *A. brassicae* treated samples of *C. sativa* at 6hr and 12 hrs after treatment. The lack of significant variation in the gene expression pattern of *ABI5* and *PYR1* genes clarifies that the receptor PYR1 and ABI5 exhibit a much more complex regulation of gene expression which may be governed by many other factors outside the scope of this study. The expression pattern of such genes which are a part of a highly intricate regulation system has to be evaluated using mutant lines.

**Figure 7:**
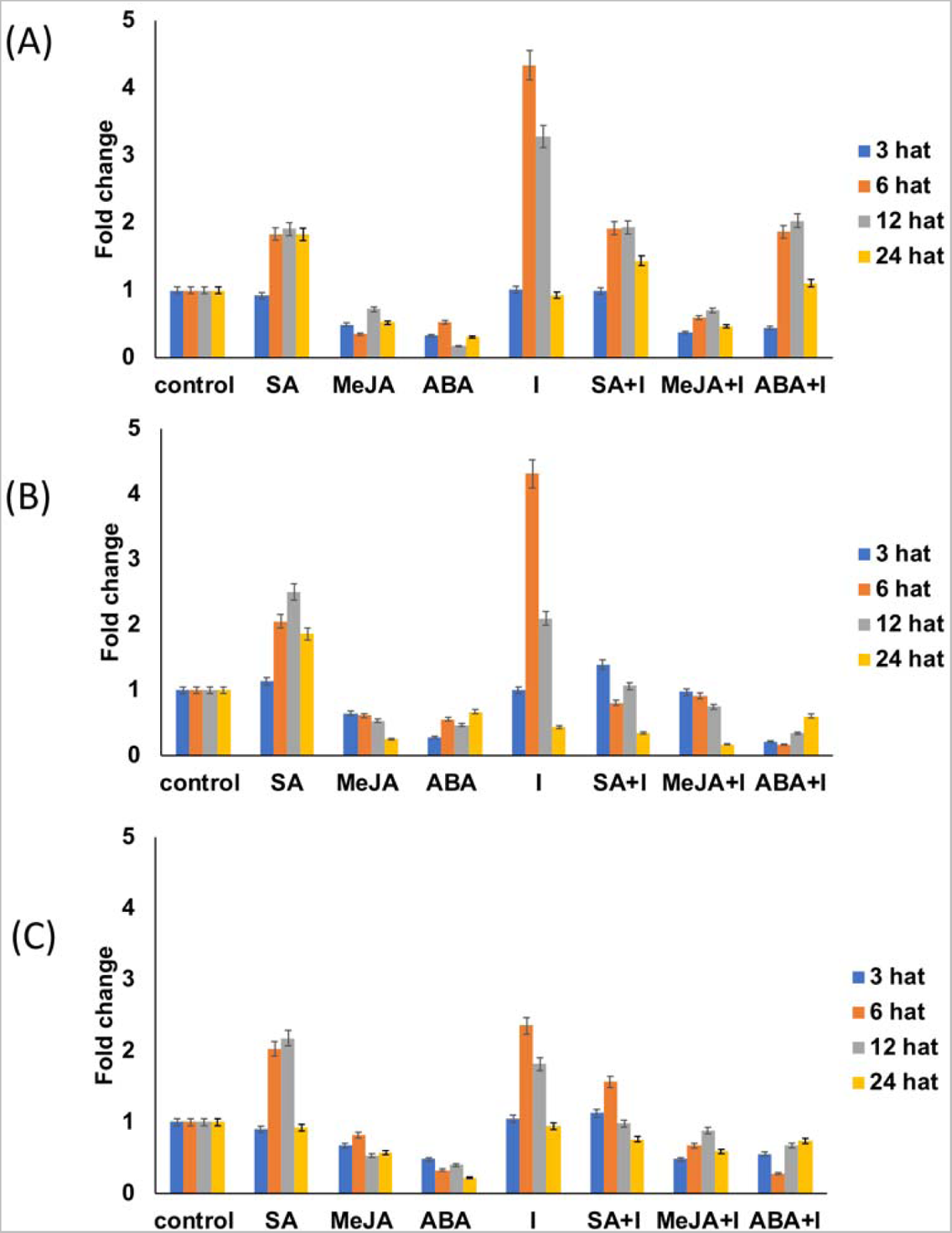
**Expression profile of *NCED3* gene in all three genotypes.** Gene expression levels of *NCED3* gene in response to various treatments at 3, 6,12 and 24 hrs after treatment (hat) in (A) *B. juncea* (B) *S. alba* (C) *C. sativa.* In this figure the treatments are- salicylic acid (SA), methyl jasmonate (MeJA), abscisic acid (ABA), I (Infection, *A. brassicae* spore suspension was used in this treatment), SA+I (SA pre-treatment followed by *A. brassicae* inoculation), MeJA+I (MeJA pre-treatment followed by *A. brassicae* inoculation) and ABA+I (ABA pre-treatment followed by *A. brassicae* inoculation).

#### 4.2.5 ABA quantification

Based on the gene expression profile of ABA biosynthesis genes and its correlation with disease progression data, the expression pattern of JA pathway and cellular JA levels, we concluded that the during the early stages of pathogen infestation the susceptible genotype failed to execute a potential JA induction giving a chance to the pathogen to hijack the signalling mechanism and use it to its advantage. In our investigation, *B. juncea* showed higher expression of ABA biosynthesis genes and lower JA accumulation in response to *A. brassicae* as compared to the tolerant genotypes. ABA-SA interaction has been found synergistic in our study. To further support our results, we performed ABA quantification in leaf tissue of all three genotypes using HPLC analysis.

In susceptible genotype *B. juncea*, the cellular ABA concentration raised after SA, *A. brassicae*, SA+I and ABA+I treatments where the highest level of induction was noticed at 12 hat (Figure 8A). *A. brassicae* and SA+I treatments cause a much higher accumulation of ABA as compared to SA treatment alone. JA treated samples show significantly low ABA concentration in the leaf tissues. These results confirmed that SA treatment induced ABA accumulation in leaf tissues and *A. brassicae* is influencing the hormonal signalling pathway in *B. juncea* by increasing ABA production to antagonize the JA mediated response and support disease establishment. In *S. alba and C. sativa*, the SA and SA+I treatment significantly raised the ABA levels in leaf samples (Figure 8B and C) interestingly, no significant accumulation of ABA in response to *A. brassicae* inoculation was noticed in the resistant genotypes. JA treatment again lowered the ABA accumulation in both the resistant genotypes. Based on gene expression analysis and hormone quantification of ABA and JA, we can conclude that ABA and JA are indeed antagonistic in the orchestra of defence signalling involving *Brassica-A. brassicae* system. The tolerant genotypes maintain lower ABA concentration after pathogen attack which did not activated ABA-SA synergy leading to successful JA mediated defence responses in these genotypes. SA supports the ABA biosynthesis and SA and ABA treated samples exhibit intense disease progression and damage. Further, we need to understand the effect of ABA treatment on the expression of SA biosynthesis pathway genes and cellular SA content to clearly understand the SA-JA-ABA interplay.

**Figure 8:**
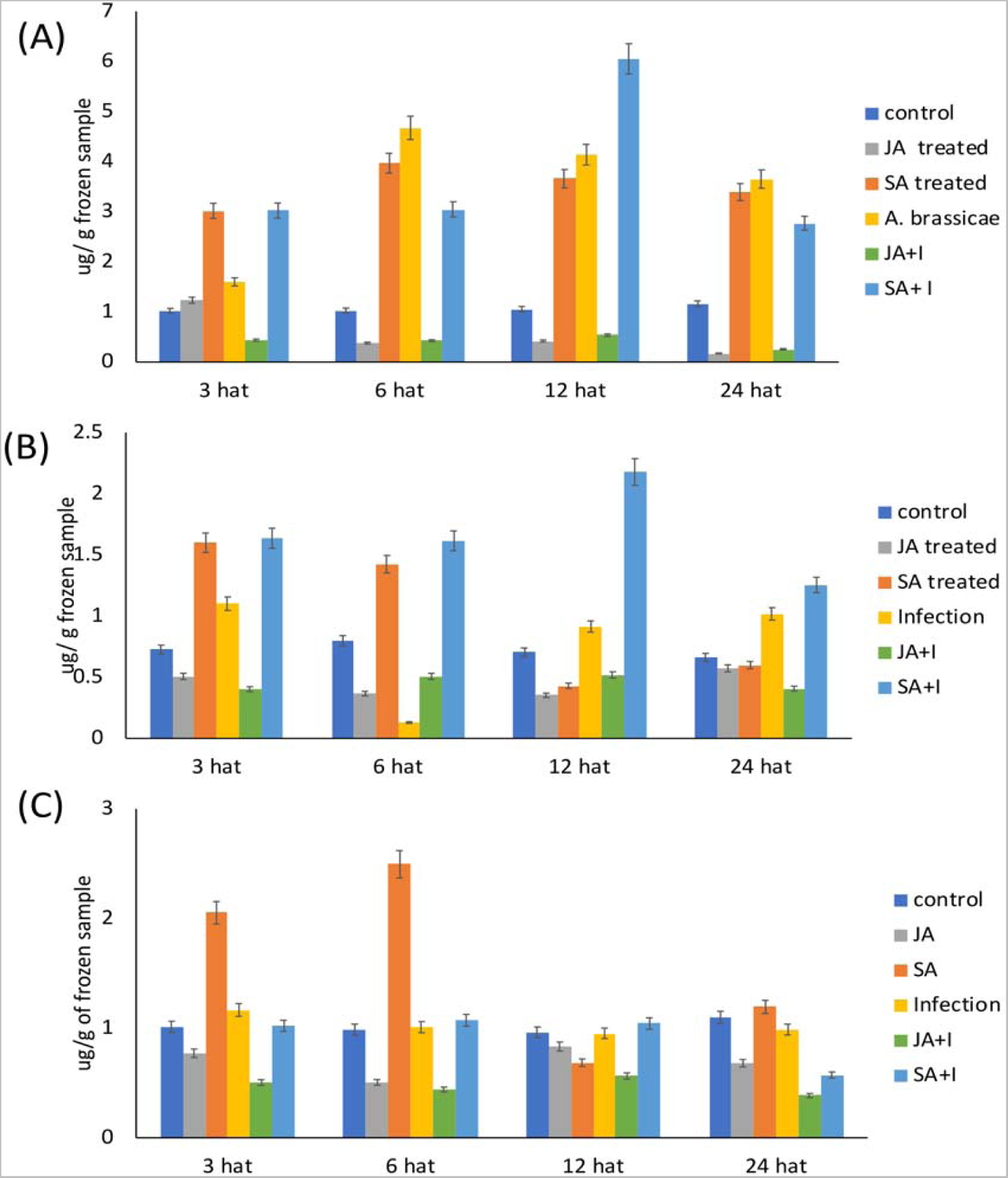
ABA levels estimated in leaf samples of all three genotypes. ABA concentration estimated in per g leaf tissue from 3 hrs to 24 hrs after treatment in (A) *B. juncea* (B) *S. alba*. (C) *C. sativa.* The standard curve was generated for quantitative estimation based on the gradient concentration of standard solution and the quantitative estimated in treated samples were calculated based on the value obtained from the mock samples of each genotype.

#### 4.2.6 Gene expression profile of SA biosynthesis pathway genes and downstream defence response gene

Salicylic acid (SA) is a phenolic compound with plant hormone activity and is recognized as an important endogenous signalling molecule in plant immunity. SA is derived from the primary metabolite chorismate, by way of two major enzymatic pathways, one involving the phenylalanine ammonia lyase pathway, and another which involves a two-step process metabolized by the enzymes isochorismate synthase (ICS) (Seilaniantz et al., 2011). The lipase-like protein EDS1 a SA biosynthesis gene represents an important node acting upstream of SA in PTI against viral, bacterial, and fungal pathogens as well as in ETI initiated by a subset of R genes (Kunkel and Brooks, 2002). We studied the expression of *EDS1* gene in all three genotypes under various treatments however we did not observe any significant variation in transcript levels of *B. juncea* and *S. alba*. The expression patterns of *PR1*, the downstream responsive gene of SA signalling pathway was also analysed for all the sample sets. In response to SA treatment, *EDS1* expression induced up to 7-fold in *B. juncea* at 12 hrs after treatment (Figure 9A). Similarly, *C. sativa* also exhibited increased *EDS1* expression in response to SA treatment. ABA treatment also induced *EDS1* gene expression however, the level of expression was found much higher in *B. juncea* (4-fold) as compared to *C. sativa* (2-fold) (Figure 9B). Interestingly in response to *A. brassicae, B. juncea* showed a 4-fold induction at 6 hrs after inoculation and no such induction was noticed in the tolerant genotype *C. sativa* (Figure 9). These results confirmed that ABA treatment significantly upregulates the expression of the major SA biosynthesis gene *EDS1* in susceptible genotype *B. juncea*. However, in *C. sativa* the induction in EDS1 gene expression was found less pronounced. These results also indicate a cooperative interaction among SA-ABA pathway genes. These results were further confirmed by quantitative estimation of SA in leaf tissue of all three genotypes using HPLC analysis.

**Figure 9:**
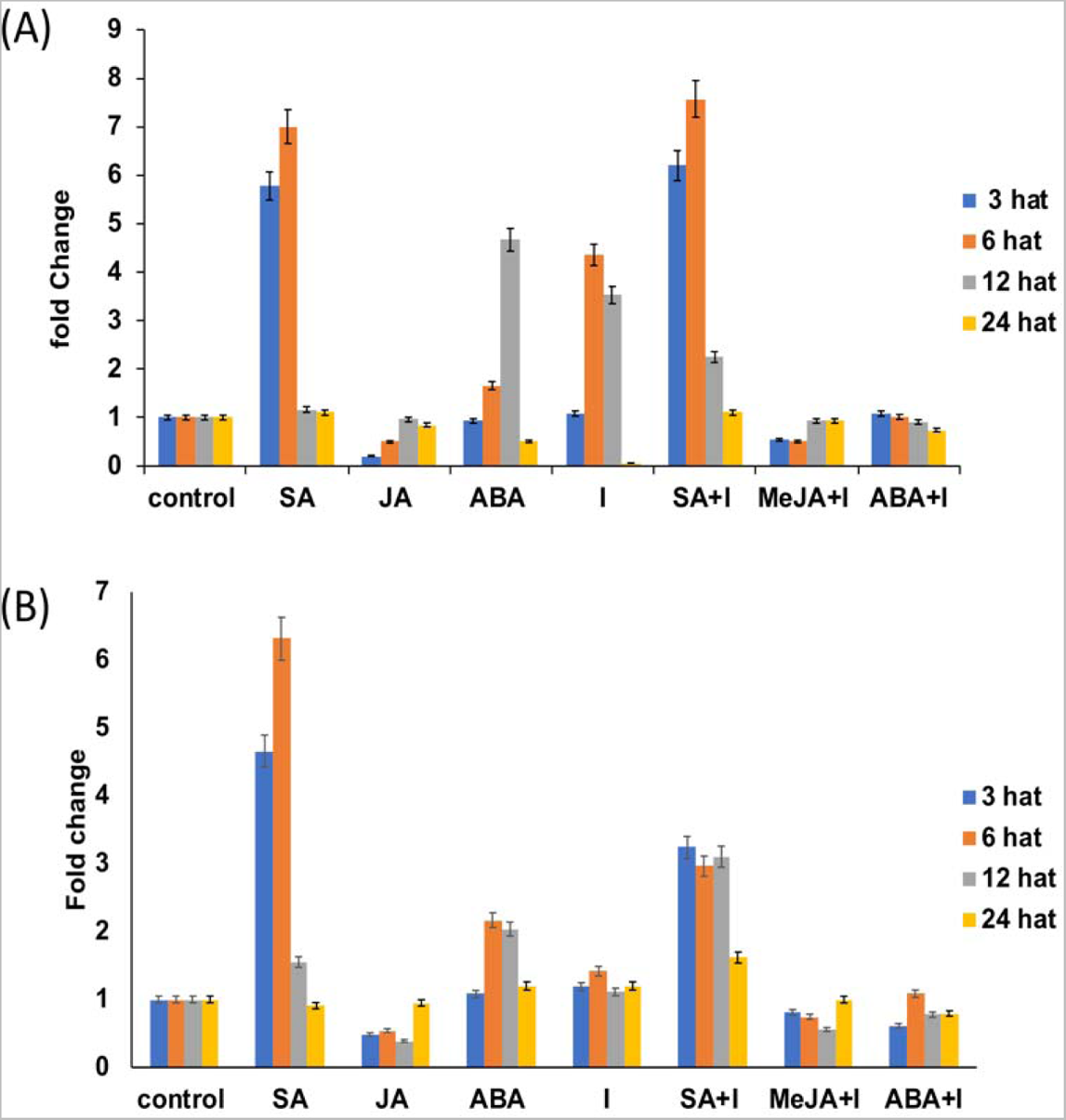
Expression profile of *EDS1* gene in all three genotypes. **Transcript levels of EDS1 gene in response to various treatments at 3, 6, 12 and 24 hrs after treatment (hat)** in (A) *B. juncea* (B) *C. sativa.* In this figure the treatments are- salicylic acid (SA), methyl jasmonate (MeJA), abscisic acid (ABA), I (Infection, *A. brassicae* spore suspension was used in this treatment), SA+I (SA pre-treatment followed by *A. brassicae* inoculation), MeJA+I (MeJA pre-treatment followed by *A. brassicae* inoculation) and ABA+I (ABA pre-treatment followed by *A. brassicae* inoculation).

PR1 gene is an important molecular indicator of systemic acquired resistance (SAR) gene in plants responsible for conferring resistance against a wide range of pathogens. Previous reports suggest that in *B. juncea* SA treatment induces the expression of PR1 and ABA treatment downregulates its expression (Ali et al., 2018). However, to the best of our knowledge, this is the first report covering the detailed analysis of cross-talk among SA-JA and ABA and its impact on hormone production in a comparative study between susceptible vs tolerant genotypes. In this investigation, an upregulation in PR1 gene expression was observed in response to SA treatment to approx. 6-fold at 3 hrs after treatment in *B. juncea* whereas, both the tolerant genotypes exhibited a lower induction of 3-fold at the same time point (Figure 10). In response to ABA treatment, *B. juncea* (susceptible) and *S. alba* (moderately tolerant) showed a non-significant induction in PR1 gene expression at 3 hrs and 6 hrs after treatment (Figures 10A and B). Whereas, *C. sativa* did not show any upregulation in response to ABA treatment (Figure 10C). The JA treatment downregulated the expression of PR1 gene in all three genotypes at all time points. In response to *A. brassicae* inoculation, an upregulation of PR1 gene up to 4-fold was noticed in *B. juncea*. This indicates signalling mechanism in *B. juncea* is biased towards SA-mediated signalling because of the inefficient JA mediated response probably caused by the manipulation of the signalling pathway due to pathogen released elicitors. In *S. alba* and *C. sativa,* no induction in PR1 gene expression was observed at given time points. This indicates a predominant JA-based signalling and defence response in the tolerant genotypes. The expression profile of PR1 gene obtained from SA+I and ABA+I further confirmed our results.

**Figure 10:**
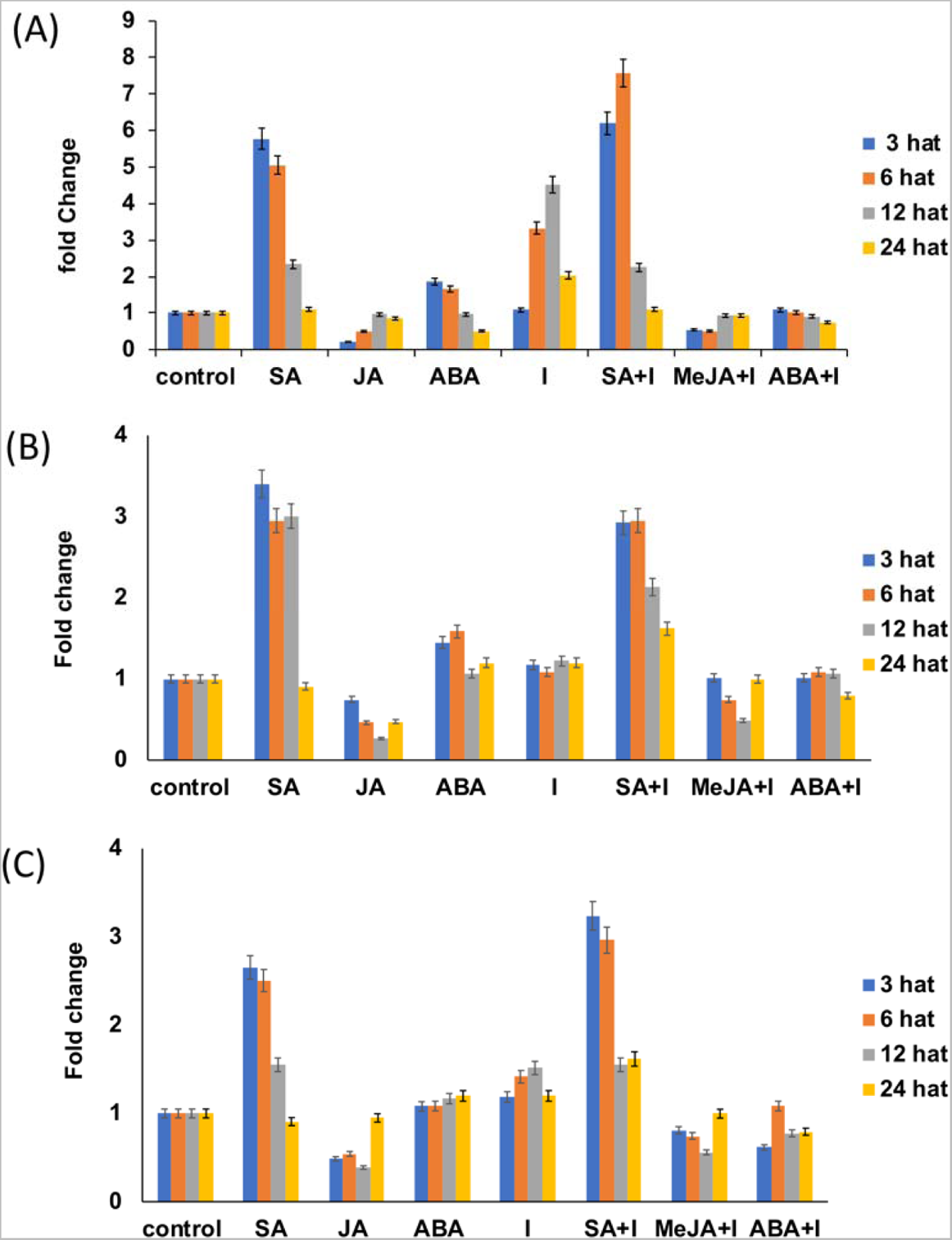
Expression profile of *PR1* gene in all three genotypes. Expression level of PR1 gene in response to various treatments at 3,6,12, and 24 hrs after treatment (hat) in (A) *B.* (B) *S. alba* (C) *C. sativa*.

#### 4.2.7 SA quantification

SA (2-hydroxybenzoic acid) is a plant signalling molecule synthesized in the chloroplast in response to the pathogen attack, it is then transported to the cytoplasm where it establishes both local and systemic acquired resistance (SAR). SA in its molecular form is toxic to cells hence it is converted into a conjugated form which is a non-toxic glycosylated form of SA and stored in vacuoles. An increasing amount of conjugated SA indicates higher SA synthesis in the plant tissue in response to pathogen attack. However, the intensity of the defence response against the invading pathogen is measured based on the amount of unconjugated SA present in the plant tissue (Mur et al., 2006).

In all three genotypes, JA and JA+I samples showed no SA accumulation (both conjugated and unconjugated) in leaf tissue at any of the time points. Graph showing lower SA concentration in leaf samples of all three genotypes after JA treatment is shown in Figure 11 and similar levels were recorded in the JA+I sample as well. ABA treatment in *B. juncea* significantly increased SA synthesis in the leaf tissue 3 hrs after treatment as evidenced by the surge recorded in conjugated SA content (3.74 μg/g) in the leaf tissue (Figure 11A). However, the unconjugated SA, a hallmark for higher signal activity was accumulated in leaf tissue from 6 hrs to 12 hrs after treatment (Figure 11A). The tolerant genotypes showed significantly lower SA concentration (both conjugated and unconjugated) as compared to the susceptible *B. juncea.* For instance, at 6 hrs after ABA treatment the conjugated SA recorded in *S. alba* and *C. sativa* was 2.82 μg/g and 2.45 μg/g respectively (Figure 11B and C). In response to *A. brassicae*, the conjugated SA content increased in the leaf tissue at 3 hrs after inoculation and a gradual fall in the conjugated SA content was recorded in the later time points of the experiment (6, 12 and 24 hrs after inoculation) (Figure 11A). However, at the same time point, a simultaneous increase in the unconjugated form of SA was recorded in the leaf tissues. This result can be explained by the phenomenon of cellular conversion of the conjugated form of SA to unconjugated form which indicates active SA mediated signalling in the leaf tissue of *B. juncea*. In *S. alba* at 3 hrs after inoculation an increase in conjugated SA was recorded up to 2.6 μg/g of leaf tissue however no increase was noticed at any time points in the unconjugated SA content (Figure 11B). This indicates that moderately tolerant *S. alba* maintains its JA-dominated defence response by restricting the conversion of the conjugated form of SA to unconjugated form in plant cells. *C. sativa* on the other hand did not exhibit any increase in cellular content of either of the SA forms explaining the JA-dominated defence response in *C. sativa* (figure 11C). The quantification of SA indicates that the signalling mechanism in the susceptible genotype *B. juncea* is dominated by the SA mediated signalling which in turn hampers the JA mediated response in *B. juncea* which is crucial for tolerance against necrotrophs. On the other hand, tolerant genotypes have evolved to quickly respond to the *A. brassicae* attack by adopting JA mediated defence response which confers tolerance against the pathogen.

**Figure 11:**
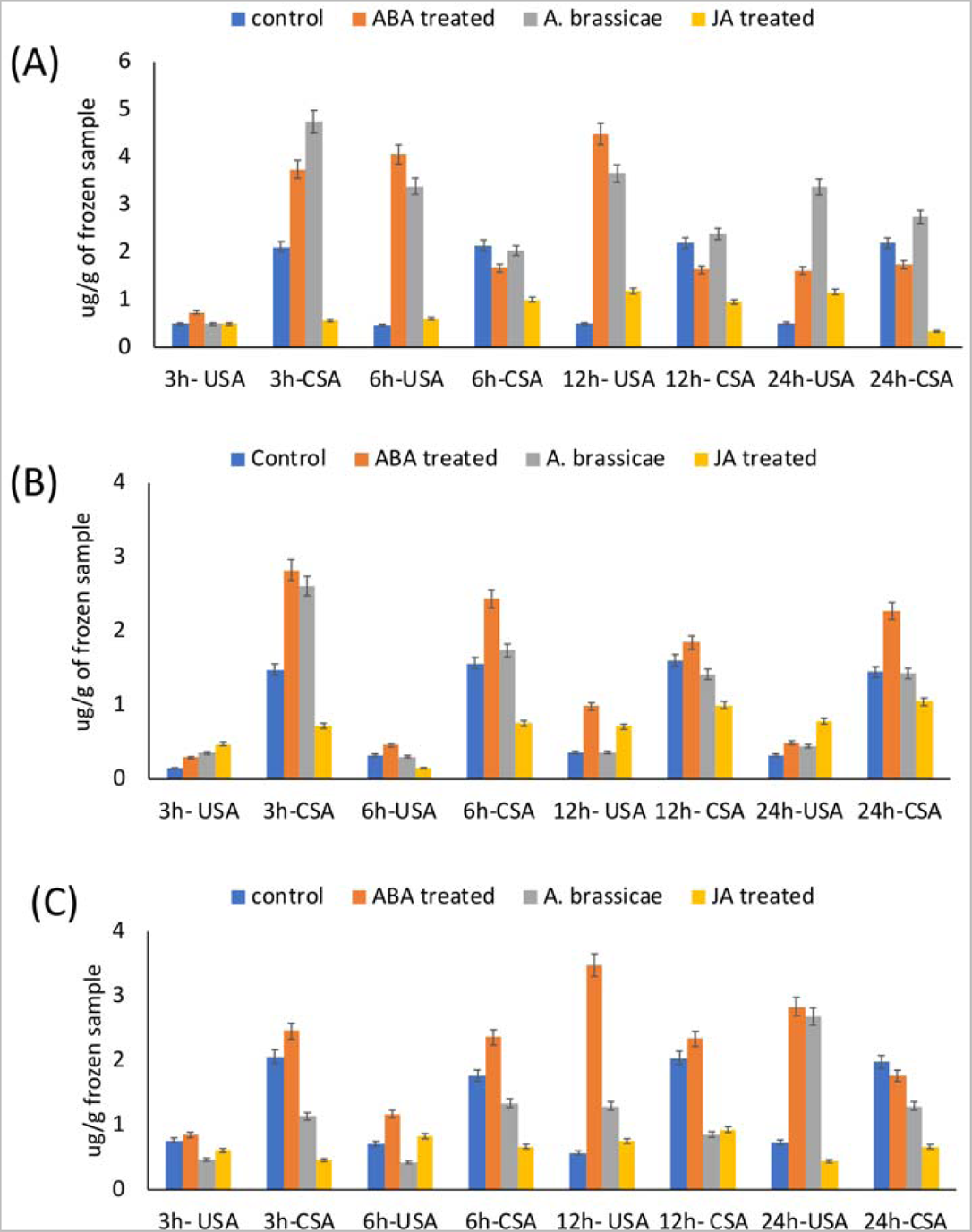
Conjugated SA and unconjugated SA levels estimated in leaf samples subjected to various treatments in all three genotypes. SA (conjugated and unconjugated) concentration estimated per g leaf tissue after various treatments from 3 hat to 24 hat in (A)B. juncea (B) *S. alba* (C) *C. sativa*. Concentration was estimated based on the area under standard curve obtained from the known gradient concentration of standard solution. In this Figure following abbreviations was used- USA-unconjugated Salicylic acid, CSA-Conjugated salicylic acid, 3h-3hrs after treatment (similar notation was used for all time points from 3hrs to 24 hrs after treatment).

## Discussion

The mustard crop is under a constant threat from numerous pathogens and *Alternaria brassicae* has been identified as one of the most pressing challenges of all (Chadar et al.,2016). The genetic constitution of any plant species contributes significantly to pathogen tolerance. Decades of breeding efforts and selection for higher yield have led to the genetic erosion of biotic stress tolerance genes in the cultivated crops rendering these crops susceptible to various deadly pathogens (Nowicki et al., 2012). However, the closely related wild genotypes of these cultivated crops are a potential reservoir of the biotic stress tolerance genes which can be utilized in breeding programs, transgenic development and genome editing programs. For this investigation, we selected three genotypes with an aim of a comparative study of molecular and physiological responses to gain insights into phytohormone cross-talk mechanisms in susceptible/tolerant genotypes. In plant defence response, phytohormones mainly SA and JA can be designated as “signal signature” which contributes to the specificity of the plant’s primary induced defence response (De Vos et al., 2005). In most plants, SA and JA trigger defence against fungal biotrophic and necrotrophic pathogens, respectively, in an antagonistic manner. It is interesting to note that SA or JA-dependent signalling pathways are not always activated exclusively on the occasion of biotrophic or necrotrophic pathogen attack (Spoel et al., 2007). For instance, a biotrophic bacterial pathogen *pseudomonas syringae* can simultaneously trigger SA and JA pathways. Pathogen being a biotroph naturally triggers the SA mediated response in plants leading to higher expression of SA responsive PR genes conferring resistance. However, the pathogen produces a JA-mimicking phytotoxin which tricks the plants to induce a JA mediated defence response which in turn antagonizes the essential SA mediated defence responses and renders the plant susceptible (Spoel et al., 2003b). In our previous study, we found that in response to a highly virulent pathotype of *A. brassicae* the susceptible genotype *B. juncea* adopts a SA mediated defence strategy which leads to higher H_2_O_2_ production in leaf tissues which works in the favour of necrotrophic pathogen *A. brassicae* and lead to higher disease progression. Whereas, the tolerant genotypes *C. sativa* and *S. alba* protect the signalling machinery from the pathogen by executing a strong and early JA induction rendering the SA-mediated signalling weak (Dixit et al., 2020b). Therefore, it can be understood that plants being exposed to thousands of pathogens in their lifetime maintains a critical balance between the SA-JA cross-talk and depending on the situation they adjust the response in favour of the most effective pathway. Virulent pathogen uses mimicking compounds of other hormones like ABA to tipoff the signalling balance of plants to work the defence response in their favour, but the tolerant genotypes evade the hijack attempt of the pathogen and exhibit tolerance against the pathogens (Seilaniantz et al., 2011). Therefore, In this investigation, we focused on deciphering the existing variation in SA-JA-ABA cross-talk in susceptible and tolerant genotypes.

In our study, we found that SA and ABA pre-treatment followed by *A. brassicae* inoculation causes aggravated disease symptoms and JA pre-treatment regulates the disease spread. However, the effect was more intense in *B. juncea* as compared to the tolerant genotypes, *S. alba* and *C. sativa* in terms of lesion size as well as overall leaf tissue damage (Figures1 and 2). These results indicated that SA-ABA are supporting the pathogen infestation whereas, JA- pre-treatment restricts the pathogen. JA pre-treatment leads to a higher cellular JA concentration which shifts the signalling mechanism towards JA mediated defence pathway resulting in tolerance against the pathogen. In a similar study, it was found that *A. brassicicola* infection in mustard leads to the accumulation of SA and enhanced expression of the SA marker gene, PR1, whereas the resistant host responded to the pathogen by upregulating the expression of JA responsive gene, *PDF1.2* (Mazumder et al., 2013). We evaluated the response of phytohormone, *A. brassicae* and a combination treatment over the hormone biosynthesis pathway centric to SA, JA and ABA in susceptible and tolerant genotypes to understand the variation present in SA-JA-ABA cross-talk.

Based on our findings we shall discuss the signalling cross-talk among SA-JA, JA-ABA and SA-ABA in terms of expression of hormone biosynthesis genes, downstream defence responsive genes and hormone quantification in each genotype. In our investigation all the genotypes exhibited an antagonistic interaction between SA-JA as evident by the downregulation of *LOX2*, *AOC2* and *PDF1.2* gene expression and low JA content in SA treated samples. Although, no surge in JA content was noticed in the SA treated leaf tissue sample of tolerant genotype but the overall JA content of *S. alba* and *C. sativa* was higher than that of the *B. juncea* at the same time point after treatment. This is because both the tolerant genotypes exhibit higher initial levels of JA before the treatment, the JA content in the untreated samples of *B. juncea* ranges from 8-10μg/g leaf tissue and the tolerant genotypes range from 15-20 μg/g leaf tissue. The residual JA in the leaf tissue of tolerant genotypes accounts for the slow deterioration of leaf tissue after SA pre-treatment followed by *A. brassicae* inoculation. In response to *A. brassicae* inoculation, both the tolerant genotypes were successful in quick activation of the JA biosynthesis pathway, as evident by the higher *AOC2* expression in *S. alba* and *C. sativa* after *A. brassicae* inoculation. This leads to higher JA accumulation in leaf tissues which quickly deploy JA mediated defence response (higher *PDF1.2* expression in tolerant genotypes) conferring resistance against *A. brassicae*. The pathogen however successfully hijacks the signalling machinery of susceptible *B. juncea* and a shift towards SA mediated defence response was noticed which is useless against *A. brassicae*. *B. juncea* inoculated samples showed induction in *EDS1* gene and PR1 gene expression and an elevated SA content in the leaf tissue confirming the activation of predominant SA mediated defence response in *B. juncea*. Hence we conclude that the tolerance towards *A. brassicae* is a result of a closely-knit signalling connection between defence response and early induction of the biosynthesis of the correct phytohormone. In the *A. brassicae-Brassica* phytosystem, the domination of JA mediated response is required to attain tolerance against the pathogen. The susceptible genotype *B. juncea* exhibits a weaker JA synthesis induction due to the predominant SA mediated response resulting in susceptibility.

In our study, we found an antagonistic interaction between ABA and JA in all three genotypes as evident by the downregulation of JA biosynthesis genes (*LOX2* and *AOC2*) and *PDF1.2* gene expression in response to ABA treatment in all three genotypes. Lower JA levels in ABA treated samples further confirmed that ABA act as a negative regulator of JA biosynthesis and JA mediated defence response. In response to *A. brassicae*, induction in ABA biosynthesis gene *NCED3* was noticed in *B. juncea* and *S. alba* and insignificant induction was noticed in tolerant genotype *C. sativa*. Further confirming the ABA levels in the *A. brassicae* inoculated samples and JA+I samples we concluded that antagonistic interaction between ABA-JA is supressing the JA induced defence response in susceptible *B. juncea* but in tolerant genotypes due to higher JA accumulation, biosynthesis of ABA is negatively regulated. However, based on the gene expression analysis it is clear that the effect of *A. brassicae* on ABA biosynthesis is complex and may involve some other TF’s and receptors which need further elaboration and confirmation.

In this investigation, the interaction between SA-ABA is synergistic. In *B. juncea*, the induction in *NCED3* gene and higher ABA accumulation in SA and SA+I treated samples endorses the synergistic relationship between ABA-SA. Further, the upregulation in *EDS1* and *PR1* in response to ABA treatment in *B. juncea* confirms that ABA induces the SA biosynthesis and hence the trade-off between SA-JA signalling is shifted towards SA mediated defence response. On comparing the expression patterns of PR1 and PDF1.2 in all the three genotypes we noticed that PR1 expression was dominating in *B. juncea* whereas, in *C. sativa* and *S. alba PDF1.2* gene expression is predominant. As it is known that inducers of these genes SA and JA work antagonistically, it can be said that response to *A. brassicae* in *S. alba* and *C. sativa* is dominated by JA induced pathway whereas, in *B. juncea* high level of SA causes an antagonistic effect. It is also likely that the pathogen *A. brassicae* is producing an ABA-mimicking toxin to trick the plant to adopt SA mediated defence response that works in the favour of the pathogen. In a similar study, it was observed that *B. cinerea* produces ABA that perturbs the host response pathway by promoting senescence and cell death (Gimenez-ibanez et al., 2016). It has been reported that few necrotrophs can release phytohormone mimics to manipulate the plant signalling machinery to its advantage (Penninckx et al., 1996). Therefore, we concluded that *A. brassicae* successfully hijacks the signalling mechanism in susceptible *B. juncea* and causes disturbance in cellular phytohormone balance by activating the ABA-SA backchannel and inducing the SA biosynthesis which deploys SA-mediated defence response (PR1 gene expression) which is unable to impart tolerance against *A. brassicae*. ABA-SA synergy suppress the JA responses leading to higher disease intensity ABA also supports the pathogen by increasing senescence and cell death in plants which led to the susceptibility in *B. juncea* (Figure 12). Tolerant genotypes evades the pathogen manipulation and exhibit an early and strong JA-mediated response against *A. brassicae*. As explained in figure 12, Early JA biosynthesis induction and accumulation supress the SA induction which led to tolerance against *A. brassicae*.

**Figure 12:**
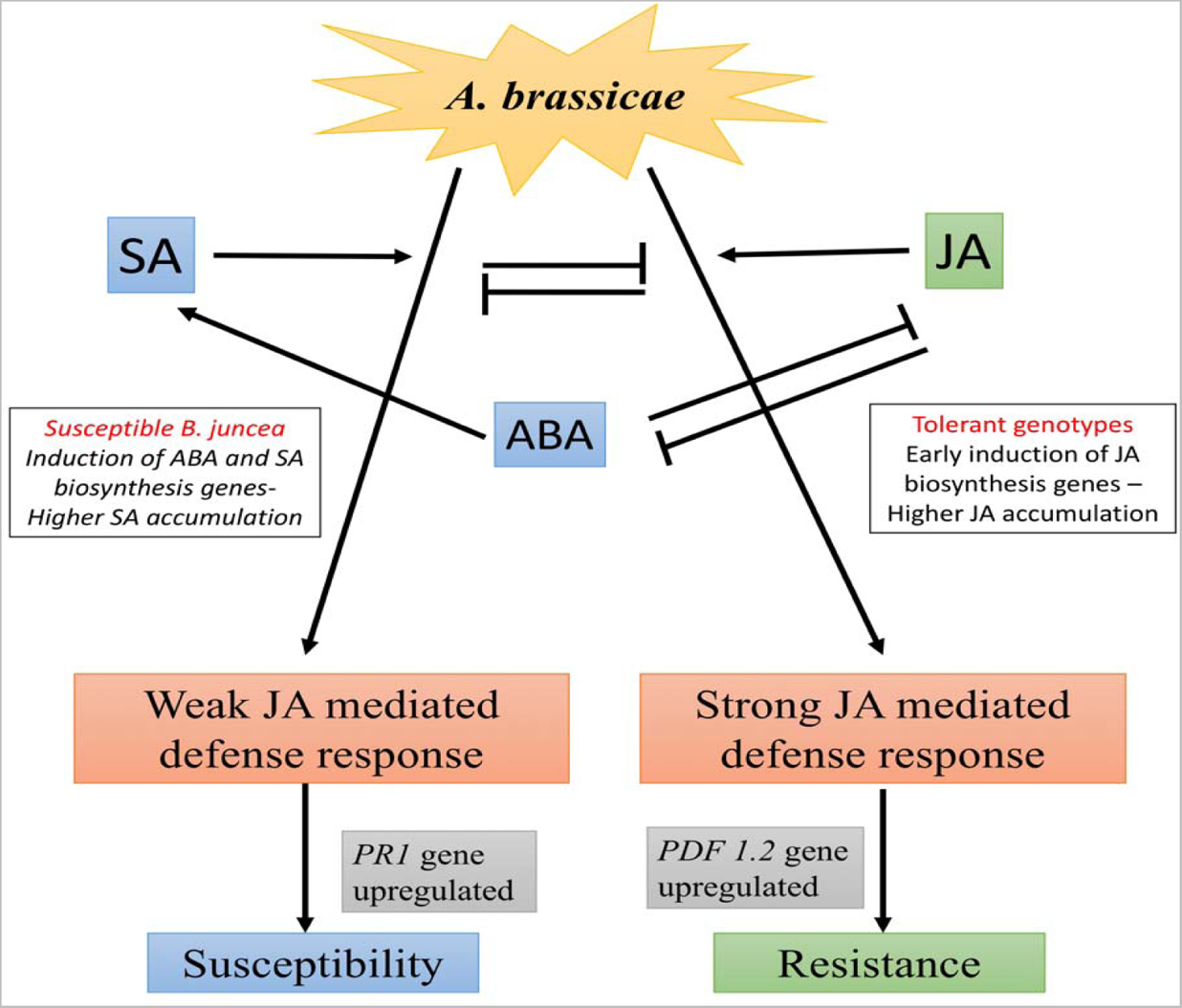
Outline of SA,JA and ABA cross-talk in susceptible B. juncea and its tolerant wild relatives. In response to *A. brassicae,* in susceptible *B. juncea* induction of ABA and SA biosynthesis and cellular accumulation of these hormones shifts the defence signalling of the plant towards SA-mediated defence response executed by higher expression of PR1 gene which fails to impart tolerance against *A. brassicae*. Tolerant genotypes execute efficient JA biosynthesis induction leading to higher JA accumulation in plant cells which lead to suppression of SA-mediated defence response and a strong induction of JA-mediated upregulation of *PDF1.2* gene resulting in tolerance against the necrotroph *A. brassicae*.

## Acknowledgments

The National Institute of Plant Biotechnology (NIPB) is acknowledged for providing research facilities. We are also thankful to IARI and National Phytotron facility for providing research facilities and infrastructure. We are thankful to the Division of plant physiology for providing the HPLC facility and to the Advanced Instrumentation Research Facility (AIRF), JNU, New Delhi for providing support and facility for GC-MS analysis.

## References

1. Ali, S., Mir, Z.A., Bhat, J.A., Tyagi, A., Chandrashekar, N., et al., 2018. Isolation and characterization of systemic acquired resistance marker gene PR1 and its promoter from *Brassica juncea*. 3Biotech. 8. https://doi.org/10.1007/s13205-017-1027-8.

2. An, C., Mou, Z., 2011. Salicylic acid and its function in plant mmunity. J. Integr. Plant Biol. 53, 412–428. https://doi.org/10.1111/j.1744-7909.2011.01043.x

3. Anderson, J.P., Badruzsaufari, E., Schenk, P.M., Manners, J.M., Desmond, O.J., et al., 2004. Antagonistic interaction between abscisic acid and jasmonate-ethylene signaling pathways modulates defense gene expression and disease resistance in *Arabidopsis*. Plant Cell 16, 3460–3479. https://doi.org/10.1105/tpc.104.025833

4. Balbi, V., Devoto, A., 2008. Jasmonate signalling network in Arabidopsis thaliana: Crucial regulatory nodes and new physiological scenarios. New Phytol. 177, 301–318. https://doi.org/10.1111/j.1469-8137.2007.02292.x.

5. Chadar, L.K., Singh, R.P., Singh, R.K., Yadav, R.R., Mishra, M.K., et al. 2016. Studies on Alternaria blight of rapeseed-mustard (*Brassica Juncea* L.) caused by *Alternaria brassicae* (Berk.) Sacc. and its integrated management. Plant Arch. 16, 897–901.

6. Clarke, J.D., Volko, S.M., Ledford, H., Ausubel, F.M., Dong, X., 2000. Roles of Salicylic Acid, Jasmonic Acid, and Ethylene in cpr-Induced Resistance in Arabidopsis. Society 12, 2175–2190. https://doi.org/10.1105/tpc.12.11.2175.

7. de Vleesschauwer, D., Yang, Y., Cruz, C.V., Höfte, M., 2010. Abscisic acid-induced resistance against the brown spot pathogen *Cochliobolus miyabeanus* in rice involves MAP kinase-mediated repression of ethylene signaling. Plant Physiol. 152, 2036–2052. https://doi.org/10.1104/pp.109.152702

8. De Vos, M., Van Oosten, V.R., Van Poecke, R.M.P., Van Pelt, J.A., Pozo, et al., 2005. Signal signature and transcriptome changes of *Arabidopsis* during pathogen and insect Attack. Mol. Plant-Microbe Interact. 18, 923–937. https://doi.org/10.1094/mpmi-18-0923

9. Derksen, H., Rampitsch, C., Daayf, F., 2013. Signaling cross-talk in plant disease resistance. Plant Sci. 207, 79–87.

10. Ding, Y., Dommel, M., Mou, Z., 2016. Abscisic acid promotes proteasome-mediated degradation of the transcription coactivator NPR1 in *Arabidopsis thaliana*. Plant J. 86, 20–34. https://doi.org/10.1111/tpj.13141

11. Dixit, S., Jangid, V.K., Grover, A., 2020a. Evaluation of stable reference genes in white mustard (*Sinapis alba*) for qRT-PCR analysis under various stress conditions. Indian J. Agric. Sci. 90, 189–194.

12. Dixit, S., Jangid, V.K., Grover, A., 2020b. Evaluation of physiological and molecular effect of variable virulence of Alternaria brassicae isolates in Brassica juncea, Sinapis alba and Camelina sativa. Plant Physiol. Biochem. 155, 626–636. https://doi.org/10.1016/j.plaphy.2020.08.025

13. Dixit, S., Jangid, V.K., Grover, A., 2019. Evaluation of suitable reference genes in Brassica juncea and its wild relative Camelina sativa for qRT-PCR analysis under various stress conditions. PLoS One 14. https://doi.org/10.1371/journal.pone.0222530

14. Fan, J., Hill, L., Crooks, C., Doerner, P., Lamb, C., 2009. Abscisic acid has a key role in modulating diverse plant-pathogen interactions. Plant Physiol. 150, 1750–1761. https://doi.org/10.1104/pp.109.137943

15. Flors, V., Ton, J., Van Doorn, R., Jakab, G., García-Agustín, P et al., 2008. Interplay between JA, SA and ABA signalling during basal and induced resistance against *Pseudomonas syringae* and *Alternaria brassicicola*. Plant J. 54, 81–92. https://doi.org/10.1111/j.1365-313X.2007.03397.x

16. Giri, P., Tasleem, M., Taj, G., Mal, R., Kumar, A., 2014. Morphological, cultural, pathogenic and molecular variability amongst Indian mustard isolates of *Alternaria brassicae* in Uttarakhand. Afr. J. Biotechnol. 13(3), 441–448. https://doi.org/10.5897/AJB2013.13198

17. Glazebrook, J., 2005. Contrasting Mechanisms of Defense Against Biotrophic and Necrotrophic Pathogens. Annu. Rev. Phytopathol. 43, 205–227.

18. Gimenez-ibanez, S., Chini, A., Solano, R., 2016. How microbes twist jasmonate signaling around their little fingers. Plants. 5(9). https://doi.org/10.3390/plants5010009

19. Audenaert, K., De Meyer G.B., Hofte M.M., 2017. Abscisic acid determines basal susceptibility of tomato to Botrytis cinerea and suppresses salicylic acid-dependent signaling mechanisms. 128, 491–501. https://doi.org/10.1104/pp.010605.1

20. Kunkel, B.N., Brooks, D.M., 2002. Cross talk between signaling pathways in pathogen defense. Curr. Opin. Plant Biol. 5, 325–331.

21. Li, X., Zhang, Y., Clarke, J.D., Li, Y., Dong, X., 1999. Identification and cloning of a negative regulator of systemic acquired resistance, SNI1, through a screen for suppressors of npr1-1. Cell 98, 329–339. https://doi.org/10.1016/S0092-8674(00)81962-5

22. Liu, X., Yang, Y.L., Lin, W.H., Tong, J.H., Huang, Z.G., et al. 2010. Determination of both jasmonic acid and methyl jasmonate in plant samples by liquid chromatography tandem mass spectrometry. Chinese Sci. Bull. 55, 2231–2235. https://doi.org/10.1007/s11434-010-3194-4

23. Mazumder, M., Das, S., Saha, U., Chatterjee, M., Bannerjee, K. et al., 2013. Salicylic acid-mediated establishment of the compatibility between *Alternaria brassicicola* and *Brassica juncea* is mitigated by abscisic acid in *Sinapis alba*. Plant Physiol. Biochem. 70, 43–51. https://doi.org/10.1016/j.plaphy.2013.04.025

24. Meena, P.D., Meena, R.L., Chattopadhyay, C., Kumar, A., 2004. Identification of critical stage for disease development and biocontrol of Alternaria Blight of Indian mustard (*Brassica juncea*). J. Phytopathol. 152, 204–209. https://doi.org/10.1111/j.1439-0434.2004.00828.x

25. Meng, X., Zhang, S., 2013. MAPK cascades in plant disease resistance signaling. Annu. Rev. Phytopathol. 51, 245–266.

26. Mur, L.A.J., Kenton, P., Atzorn, R., Miersch, O., Wasternack, C., 2006. The outcomes of concentration-specific interactions between salicylate and jasmonate signaling include synergy, antagonism, and oxidative stress leading to cell death.Plant Physiol. 140, 249–262. https://doi.org/10.1104/pp.105.072348.

27. Nakurte, I., Keisa, A., Rostoks, N., 2012. Development and validation of a reversed-phase liquid chromatography ethod for the simultaneous determination of indole-3-acetic acid, indole-3-pyruvic acid, and abscisic acid in barley (*Hordeum vulgare* L.). J. Anal. Methods Chem. 2012, 1–6. https://doi.org/10.1155/2012/103575

28. Nowicki, M., Nowakowska, M., Niezgoda, A., Kozik, E., 2012. Alternaria black spot of crucifers: Symptoms, importance of disease, and perspectives of resistance breeding. Veg. Crop. Res. Bull. 76, 5–19. https://doi.org/10.2478/v10032-012-0001-6

29. Pandey, D., Raj, S., Kunnumal, C., Gaur, M., 2016. Plant defense signaling and responses against necrotrophic fungal pathogens. J. Plant Growth Regul. 35, 1159–1174. https://doi.org/10.1007/s00344-016-9600-7

30. Peña-Cortés, H., Barrios, P., Dorta, F., Polanco, V., Sánchez, C. et al., 2004. Involvement of jasmonic acid and derivatives in plant response to pathogen and insects and in fruit ripening. J. Plant Growth Regul. 23, 246–260. https://doi.org/10.1007/BF02637265

31. Penninckx, A.M.A., Eggermont, K., Terras, F.R.G., Bart, P., 1996. Pathogen-induced systemic activation of a plant defensin gene in *Arabidopsis* follows a salicylic acid-independent pathway. The Plant Cell. 8, 2309–2323.

32. Penninckx, I.A.M.A., Thomma, B.P.H.J., Buchala, A., Métraux, J., Broekaert, W.F., 1998. Concomitant activation of jasmonate and ethylene response pathways is required for induction of a plant defensin gene in *Arabidopsis*. The Plant Cell.10, 2103–2113.

33. Pieterse, Corné M.J., Van der Does, D., Zamioudis, C., Leon-Reyes, A., Van Wees, S.C.M., 2012. Hormonal modulation of plant Immunity. Annu. Rev. Cell Dev. Biol. 28, 489–521.

34. Pieterse, M.J., 2008. Cross-talk in defense signaling. 146, 839–844. https://doi.org/10.1104/pp.107.112029

35. Qin, X., Zeevaart, J.A.D., 2002. Overexpression of a 9-cis-Epoxycarotenoid Dioxygenase Gene increases abscisic acid and phaseic acid levels and enhances drought tolerance. Plant physiol. 128(2), 544–551. https://doi.org/10.1104/pp.010663.

36. Rai, S.K., Charak, D., Bharat, R., 2016. Scenario of oilseed crops across the globe. Plant Arch. 16, 125–132.

37. Rowe, H.C., Walley, J.W., Corwin, J., Chan, E.K., Dehesh, K. et al., 2010. Deficiencies in jasmonate-mediated plant defense reveal quantitative variation in Botrytis cinerea pathogenesis. PLoS Pathog 6. e1000861. https://doi.org/10.1371/journal.ppat.1000861

38. Savary, S., Teng, P.S., Willocquet, L., Nutter, F.W., 2006. Quantification and modeling of crop losses : A review of purposes. Annu. Rev. Phytopathol 44, 89–112.

39. Sharma, G., Kumar, V.D., Haque, A., Bhat, S.R., Prakash, S. et al., 2002. Brassica coenospecies: A rich reservoir for genetic resistance to leaf spot caused by Alternaria brassicae. Euphytica 125, 411–417. https://doi.org/10.1023/A:1016050631673

40. Seilaniantz, A.R., Grant, M., Jones, J.D.G., 2011. Hormone crosstalk in plant disease and defense: More than just Jasmonate-Salicylate antagonism. Annu. Rev. Phytopathol. 49, 317–343.

41. Shigenaga, A.M., Argueso, C.T., 2016. No hormone to rule them all: Interactions of plant hormones during the responses of plants to pathogens. Semin. Cell Dev. Biol. 56, 174–189. https://doi.org/10.1016/j.semcdb.2016.06.005

42. Singh, A.K., Choudhary, A.K., Kumari, A., Kumar, R., 2017. Towards oilseeds sufficiency in India: Present status and way forward. J. AgriSearch 4, 80–84.

43. Spoel, S.H., Johnson, J.S., Dong, X., 2007. Regulation of tradeoffs between plant defenses against pathogens with different lifestyles. Proc. Natl. Acad. Sci. U.S.A. 104, 18842–18847

44. Spoel, S.H., Koornneef, A., Claessens, S.M.C., Korzelius, J.P., Pelt, J.A. et al., 2003a. NPR1 Modulates Cross-Talk between Salicylate-and Jasmonate-Dependent Defense Pathways through a Novel Function in the Cytosol. Plant Cell 15, 760–770. https://doi.org/10.1105/tpc.009159

45. Spoel, S.H., Koornneef, A., Claessens, S.M.C., Korzelius, J.P., Van Pelt, J.A. et al., 2003b. NPR1 modulates cross-talk between salicylate- and jasmonate-dependent defense pathways through a novel function in the cytosol. Plant Cell. 15, 760–770. https://doi.org/10.1105/tpc.009159

46. Thomma, B.P.H.J., Nelissen, I., Eggermont, K., Broekaert, W.F., 1999. Deficiency in phytoalexin production causes enhanced susceptibility of *Arabidopsis thaliana* to the fungus *Alternaria brassicicola*. Plant J. 19, 163–171. https://doi.org/10.1046/j.1365-313X.1999.00513.x

47. Untergasser, A., Cutcutache, I., Koressaar, T., Ye, J., Faircloth, B.C., Remm, M., et al., 2012. Primer3-new capabilities and interfaces. Nucleic Acids Res. 40(15), https://doi.org/10.1093/nar/gks596

48. Wasternack, C., Song, S., 2017. Jasmonates: Biosynthesis, metabolism, and signaling by proteins activating and repressing transcription. J. Exp. Bot. 68, 1303–1321. https://doi.org/10.1093/jxb/erw443

49. Xu, J., Audenaert, K., Hofte, M., de Vleesschauwer, D., 2013. Abscisic Acid promotes susceptibility to the rice leaf blight pathogen Xanthomonas oryzae pv oryzae by suppressing salicylic acid-mediated defenses. PLoS One. 8. https://doi.org/10.1371/journal.pone.0067413

